# *APOE* genotype affects the composition of secreted apoE lipoprotein particles and cellular lipid profiles in pericytes

**DOI:** 10.64898/2026.11.16.766920

**Authors:** Evelina Voloviceva, Sandra Goetze, Safiyo Haji Abukar, Stephanie Hausler, Annika Jagels, Alaa Othman, Arnold von Eckardstein, Jérôme Robert

**Affiliations:** Institute of Clinical Chemistry, University Hospital of Zurich, University of Zurich, Zurich, Switzerland; BioMed Program, Life Science Zurich Graduate School, University of Zurich, Zurich, Switzerland; Swiss Multi-Omics Center, ETH Zurich, Zurich, Switzerland; Institute of Molecular and Translational Medicine, ETH Zurich, Zurich, Switzerland; Swiss Institute of Bioinformatics, Lausanne, Switzerland; Functional Genomics Center Zurich, ETH Zurich and University of Zurich, Zurich, Switzerland

## Abstract

Impairment of the blood-brain barrier (BBB) is often observed in Alzheimer’s disease patients, especially those who carry the *APOE4* allele of apolipoprotein E. Brain pericytes are crucial for maintaining the BBB. Yet, the mechanisms linking *APOE* genetic variation to pericyte-mediated pathologies are not fully understood. Given the importance of pericyte-secreted factors in cerebrovascular homeostasis, we hypothesized that the protein and lipid compositions of pericyte-secreted apoE lipoproteins (LpEs) would differ depending on the *APOE* genotype. Using liquid chromatography-mass spectrometry, we investigated the lipid and protein components of LpEs secreted by human induced pluripotent stem cell-derived *APOE-*isogenic pericytes.

Our results demonstrate that *APOE2/E2*, *APOE3/E3*, and *APOE4/E4* pericytes secrete LpEs with distinct lipid and protein compositions. ApoE4 particles (LpE4) have high total lipid levels, LpE3 are enriched in ether-linked phospholipids, and LpE2 in ester-linked phosphatidylcholines and short-chain sphingomyelins. Extracellular matrix proteins are highly represented in LpE4, followed by LpE2. In particular, fibronectin is abundant in LpE4 and LpE2, but not in LpE3. While cellular lipid levels and lipid subclass compositions also differ between *APOE*-isogenic pericytes, they do not mirror the lipid profiles of secreted LpE. The most differentially abundant cellular lipid subclasses include triglycerides, sphingomyelins, glycosphingolipids (Hex3Cer, Hex2Cer, HexCer), and ceramides.

In conclusion, the composition of pericyte-secreted LpEs varies by *APOE* genotype, suggesting distinct functions in brain homeostasis and neurodegenerative disease. Differences in cellular lipids further suggest *APOE*-mediated changes in lipid metabolism, highlighting the role of vascular cells in brain health and disease.

## 1. Introduction

Perivascular cells, also known as pericytes, are located along the walls of capillaries. In the central nervous system (CNS), they maintain the blood-brain-barrier (BBB), regulate microvascular blood flow, and support angiogenesis [1]. Vascular dysfunction, including BBB breakdown associated with pericyte injury, has been proposed as one of the earliest pathogenic mechanisms of Alzheimer’s disease (AD), occurring before the pathological hallmarks of AD: amyloid deposition, tau accumulation, and cognitive impairment [2,3]. The *APOE* genotype is one of the factors promoting pericyte dysfunction. Accelerated pericyte loss and BBB impairment are observed in *APOE4* carriers with AD [4], yet the mechanisms by which pericyte-derived apoE contribute to neurodegeneration remain unclear.

Several mechanisms that link genetic variation in *APOE* to pericyte-mediated CNS pathologies have been proposed. First, apoE3 and apoE4 differentially affect pericyte migration and adhesion, which are crucial for maintaining cerebrovascular homeostasis [5]. Second, *APOE4* pericytes adopt myofibroblast-like phenotypes accompanied by increased secretion of extracellular matrix (ECM) components that facilitate amyloid deposition [6]. Third, lipoproteins secreted by *APOE3* pericytes (LpE3) provide better support in barrier formation by endothelial cells compared to LpE4 [7]. Lastly, *APOE4* pericytes accumulate higher levels of the proinflammatory cytokine cyclophilin A (CypA) and matrix metalloproteinase-9 (MMP-9), which lead to MMP-9-mediated BBB breakdown [4]. These examples demonstrate that the *APOE* genotype influences pericyte functions and highlight the importance of pericyte-secreted factors, such as lipoproteins, cytokines, and ECM components in cerebrovascular health. Emerging evidence suggests that apoE lipoproteins (LpEs) contain many additional proteins and lipids beyond apoE [8–10]. Therefore, we hypothesized that the protein and lipid compositions of pericyte-secreted LpEs differ depending on the *APOE* genotype. In this study, we characterized the compositions of LpEs secreted by *APOE*-isogenic induced pluripotent stem cell (iPSC)-derived pericytes and compared these secreted profiles to the cellular lipidome and proteotype of the cells.

## 2. Methods

### 2.1 iPSC culture

*APOE*-isogenic iPSC lines (parental line BIONi037-A — genotyped *APOE3/E3*, derived from a 77-year-old healthy female donor and CRISPR/Cas9-edited lines BIONi037-A-1 — *APOE* KO, BIONi037-A-2 — *APOE2/E2*, BIONi037-A-4 — *APOE4/E4*) were purchased from The European Bank for induced pluripotent Stem Cells (EBiSC) [11,12]. The iPSCs were cultured on Matrigel-coated (354277 Corning) plates in mTeSR Plus medium (100–0276 StemCell Technologies) and passaged every 3-4 days. Culture media were replaced every 1-2 days. For passaging, cells were washed with PBS (10010023 Gibco), incubated with ReLeSR (100-0483 StemCell Technologies) for 5 min at 37°C with 5% CO_2_, detached in 1 mL culture medium, and transferred to new Matrigel-coated plates.

### 2.2 Pericyte differentiation

Pericyte differentiation through the neural crest (NC) intermediate stage was carried out according to the protocol adapted from Faal *et al.* [13], as previously described [10].

### 2.3 Pericyte marker expression analysis by immunofluorescence staining

The iPSC-derived pericytes were seeded in Matrigel-coated black 96-well optical bottom plates (165305 Thermo Scientific) (0.01 x 10^6^ cells/well). The immunofluorescence staining was carried out as previously described [10]. Briefly, the cells were fixed with 4% paraformaldehyde (PFA) and extensively washed with phosphate-buffer saline (PBS). Blocking solution containing 5% normal donkey serum and 0.2% bovine albumin (BSA) in PBS was applied for one hour. After one hour, the cells were incubated overnight at 4°C with mouse anti-PDGFRβ (RRID: AB_1269704, 1:250) and rabbit anti-NG2 (RRID: AB_2877152, 1:200) antibodies diluted in the blocking solution. Following extensive PBS washes, the cells were incubated with donkey anti-mouse AlexaFluor 647 (RRID: AB_162542, 1:500) or donkey anti-rabbit AlexaFluor 568 (RRID: AB_162542, 1:500) secondary antibodies diluted in PBS. Images were acquired using Cytation 5 cell imaging multimode reader (Agilent BioTek) at 20x magnification and processed with Gen5 software (Agilent BioTek) and Fiji ImageJ (1.54f, NIH).

### 2.4 Gene expression analysis using RT-qPCR

Total RNA was isolated from respective cells using the NucleoSpin™ Mini Kit for RNA Purification (740955 Macherey Nagel) following manufacturer’s recommendations. RNA (0.5 µg per reaction) was reverse transcribed into cDNA using RevertAid First Strand Synthesis kit (K1621 Thermo Scientific). Real-time qPCR reactions were carried out in a Light Cycler 480-II (Roche) using the LightCycler® 480 SYBR Green I Master Mix (04887352001 Roche) and respective primers for genes of interest: *GAPDH*: for: CCCATGTTCGTCATGGGTGT; rev: TGGTCATGAGTCCTTCCACGATA, *POU5F1*: for: GCAAAGCAGAAACCCTCGTG, rev: GATCTGCTGCAGTGTGGGT, *SOX2*: for: CAACGGCAGCTACAGCATGA, rev: CTCGGACTTGACCACCGAAC, *NANOG*: for: GAGATGCCTCACACGGAGAC, rev: GGGTTGTTTGCCTTTGGGAC, *SOX10*: for: ATGAACGCCTTCATGGTGTGGG, rev: CGCTTGTCACTTTCGTTCAGCAG, *PDGFRB*: for: ACAGACTCCAGGTGTCATCC, rev: AGTTGACCACCTCATTCCCG, *MCAM*: for: GCCAGTCCTCATACCAGAGC, rev: TCTTACGAGACGGGGGTAGC, *CSPG4*: for: GTCCTGCCTGTCAATGACCAAC, rev: CTGTGGGTGAGATCATGTGG.

Gene expression was quantified with the ΔΔCt method. All results were normalized to *GAPDH* as housekeeping gene. For each marker gene, expression values were normalized (set to 1) to the reference cell type: iPSCs for *POU5F1*, *SOX2* and *NANOG*, neural crest cells for *SOX10*, and pericytes for *PDGFRB*, *MCAM* and *CSPG4*.

### 2.5 Media collection for the characterization of secreted LpEs

Terminally differentiated pericytes were cultured in delipidated pericyte culture media (1201 ScienCell Pericyte Medium with growth supplements and 2% delipidated FBS). After 72 hours with no medium changes, conditioned media were collected and processed as previously described [10].

### 2.6 ApoE concentration determination

The concentrations of apoE in cell-conditioned media were determined by apoE ELISA, adapted from Fan *et al*. [14] as previously described [10].

### 2.7 Immunoprecipitation of LpEs

Antibody-bead conjugates were prepared with the Dynabeads® Antibody Coupling Kit (14311D Thermo Scientific) as previously described [10]. Unconcentrated pericyte-conditioned media were added to the antibody-bead conjugates with volumes adjusted to contain 150 ng apoE per one reaction. Final reaction volumes were equalized across all samples by adjusting sample input volumes with vehicle medium. Separate control samples contained *APOE* KO pericyte conditioned media or vehicle media without cell contact, added to antibody-bead conjugates at volumes matched to experimental samples. ApoE immunoprecipitation was carried out as previously described [10].

### 2.8 Proteotyping of LpEs

The analysis included immunoprecipitated LpEs from *APOE3/E3*, *APOE2/E2*, and *APOE4/E4* iPSC-derived pericytes. The immunoprecipitates obtained after incubating the antibody-bead conjugate with either *APOE* KO pericyte-conditioned media or vehicle media were processed in the same manner as the experimental samples. The immunoprecipitates were lysed and processed with iST 8x kits (P.O.00001 PreOmics) to obtain the peptide mix prior to analysis by liquid chromatography coupled to mass spectrometry (LC-MS/MS). Mass spectrometry and raw files analyses, as well as the curation of initial protein list were performed as previously described [10]. Briefly, peptides were separated on a Vanquish™ Neo UHPLC system using a 50 cm μPAC™ Neo column (Thermo Fisher Scientific). The immunoprecipitates were eluted at 300 nL min⁻¹ using a gradient of 8–32% mobile phase B (80% acetonitrile, 0.1% formic acid) over 40 min, followed by 32–50% B over 4 min and analyzed by data-dependent acquisition on an Orbitrap Exploris™ 480. Raw files were processed in FragPipe v23.0 using the DDA workflow. MSFragger v4.3 searched spectra against the reviewed UniProt *Homo sapiens* reference proteome (June 2025), supplemented with common contaminants and reverse decoys. Searches used strict trypsin specificity, up to two missed cleavages, and 20 ppm precursor and fragment mass tolerances. Carbamidomethylation of cysteine was fixed; methionine oxidation and protein N-terminal acetylation were variable. Peptide and protein identifications were filtered with 1% FDR. Label-free quantification was performed using IonQuant with MaxLFQ and match-between-runs enabled.

Only proteins enriched (fold change (FC) ≥ 2 and adjusted *P* value < 0.05) in each experimental condition compared to *APOE* KO conditioned medium controls were defined as LpE components. Proteins enriched (FC ≥ 2 and adjusted *P* value < 0.05) in immunoprecipitates from pericyte-conditioned media (including *APOE* KO conditioned medium) compared to vehicle medium controls were defined as apoE immunoprecipitation-associated (apoE IP-associated). Protein filtering criteria and results are included in the Supplementary File 1.

### 2.9 Cell proteotyping

The iPSCs were collected after two passages post-thawing. The neural crest cells (NC) were collected on day 6 of the iPSC-to-pericyte differentiation. The iPSC-derived pericytes were collected on day 24 of the differentiation. The cells were washed with PBS, detached with Accutase and counted. Suspensions containing 1 x 10^5^ cells per replicate (well) were transferred to 2 mL Eppendorf tubes and centrifuged at 300g for 5 minutes at 4°C. Supernatants were removed and the cells were washed with PBS. Following centrifugation, the supernatants were then discarded, and cell pellets were stored at –70°C until further processing. Cell pellets were processed with iST 8x kits (P.O.00001 PreOmics) to obtain the peptide mix prior to LC-MS/MS analysis. Total cell lysates were eluted at 300 nL min⁻¹ using a gradient of 8–32% mobile phase B (80% acetonitrile, 0.1% formic acid) over 75 min, followed by 32–52% B over 10 min. MS acquisition was performed as described above. Raw files were processed in FragPipe v23.0 using the DDA+ workflow.

### 2.10 Cell collection for lipidomics

iPSC-derived pericytes were cultured until day 24 of the differentiation and cell pellets obtained as described above. Lipid extraction from the frozen cell pellets was carried out using the butanol:methanol (BuMe, 1:1, v/v) method prior to LC-MS/MS analysis. The hydrophobic protein pellets remaining after lipid extraction were dried with a stream of nitrogen, solubilized with 60 µl of 0.1 M NaOH, and incubated in a thermomixer for 10 min at 50°C and 1000 rpm. Protein concentrations were then determined using the Qubit™ Protein Assay Kit (A50669 Invitrogen). Total protein amounts were used for cellular lipidomics data normalization.

### 2.11 Lipidomics of LpEs and cells

For untargeted lipidomics, we included the same conditions as for proteotyping. Lipid extraction, lipid profiling, and data analysis were adapted from Ganguin *et al*. [15] and carried out as previously described [10].

Only lipids enriched (FC ≥ 1.5 and adjusted *P* value < 0.05) in each experimental condition compared to *APOE* KO conditioned medium controls were defined as LpE components. Lipids enriched (FC ≥ 1.5 and adjusted *P* value < 0.05) in immunoprecipitates from pericyte-conditioned media (including *APOE* KO conditioned medium) compared to vehicle medium controls were defined as apoE IP-associated. Fold change thresholds of 1.5 were applied for sample versus control comparisons to capture biologically relevant secreted and signaling lipids, where less than 2-fold changes in abundance can have functional consequences [16]. Lipid filtering criteria and results are included in the Supplementary File 2.

### 2.12 Statistics

Normality or lognormality of data distribution were assessed with the Shapiro-Wilk test. Comparisons between two groups were performed using Welch’s t-test, comparisons between multiple groups — with one-way ANOVA followed by Dunnett’s (where the control group is present) or Tukey’s (for multiple comparisons without a clear control group) post hoc tests. Mann-Whitney and Kruskal-Wallis tests were used as non-parametric alternatives for two-group and multiple group comparisons, respectively. Data visualization and statistical analyses were conducted in GraphPad Prism and R software. ChatGPT (model GPT-4o, OpenAI) and Claude (model Sonnet 5, Anthropic) Large Language Models were used for R code troubleshooting and optimization.

Lipidomics and proteomics data analysis was carried out in R using Bioconductor packages *lipidr* and *limma*. Data were visualized with *pheatmap*, *ggplot2*, and *eulerr* packages. Lipidomics data were normalized to internal standards of known concentration (for cellular lipid profiling, data were additionally normalized to total protein content) and log_2_-transformed. Relative lipid abundances were calculated from normalized linear data, before log_2_-transformation. Absolute concentrations were determined by normalizing analyte signal to internal standards of known concentration and to total protein content for cellular lipid profiling.

Proteomics data were log_2_-transformed and median-centered. Relative protein abundances were calculated from median-centered linear data, before log_2_-transformation. Proteins with at least 40% non-missing value across all samples were retained for analysis. Missing values were imputed using the minimum probability (MinProb) method from the *imputeLCMD* R package. For heatmap and volcano plot visualization, proteomics data were additionally normalized using APOE intensity after median-centering. *P* values were adjusted for multiple testing with the Benjamini-Hochberg method to control the false discovery rate (FDR). Differences were considered statistically significant at p < 0.05, FDR < 0.05, and |FC| ≥ 2 (|log2FC| ≥ 1), where applicable. For all analyses, one replicate represented one well derived from an independent differentiation. Three replicates per condition were analyzed for –omics analyses and at least three differentiation replicates per condition for qPCR.

## 3. Results

### 3.1 *APOE*-isogenic iPSCs differentiate into apoE-secreting pericytes

To investigate the compositions of LpEs secreted by pericytes with different *APOE* genotypes, we differentiated *APOE*-isogenic human iPSCs into pericytes (Fig. 1 A). To assess whether the differentiation had occurred, we analyzed the expression of pericyte markers in the resulting cells on the RNA and protein levels. We included iPSCs, intermediate-stage neural crest (NC) cells, and terminally differentiated pericytes in the gene expression analyses.

**Figure 1.**
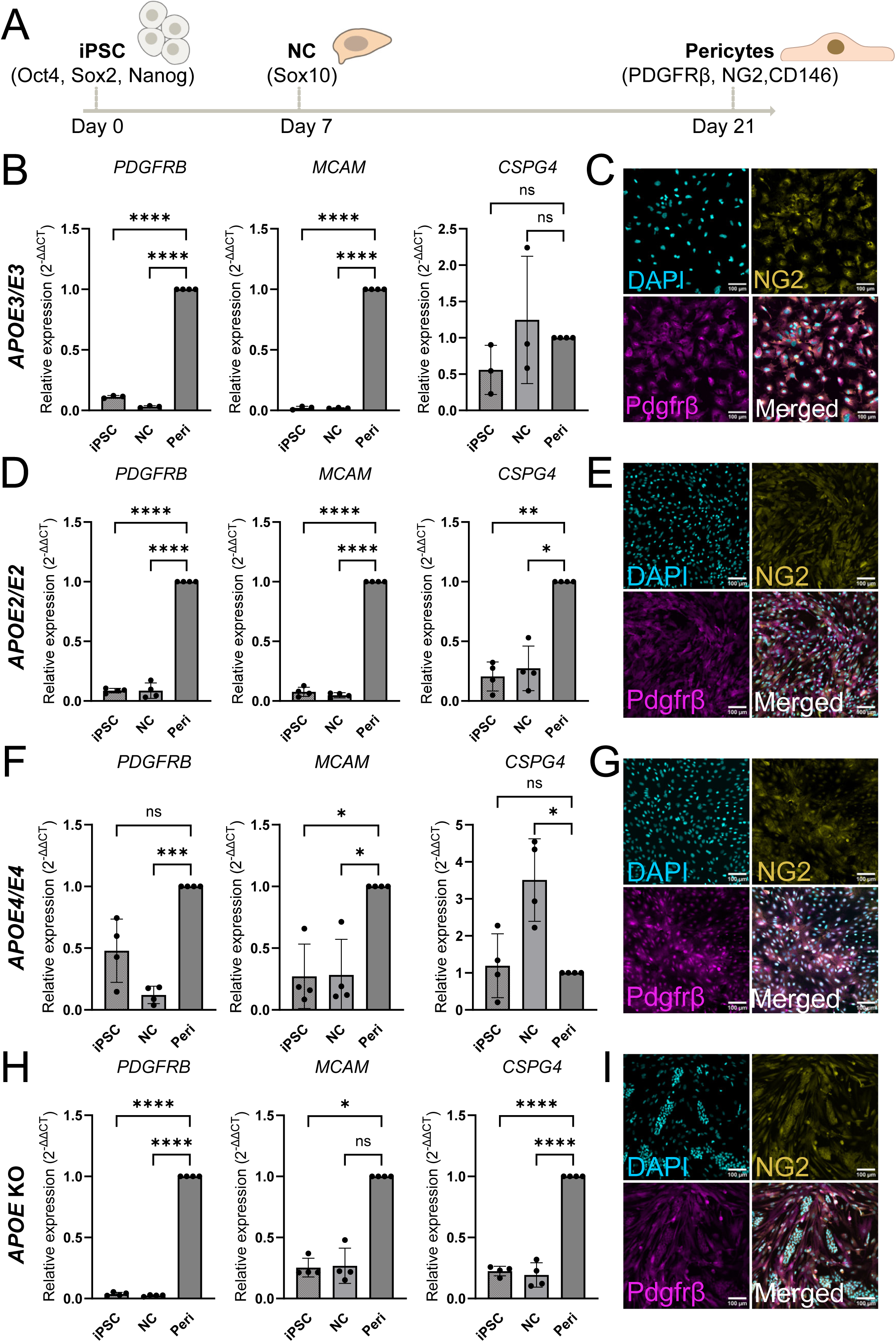
Characterization of iPSC-derived *APOE-*isogenic pericytes. A), Schematic differentiation route of iPSCs (markers: Oct4, Sox2, Nanog) to pericytes (markers: CD146, PDGFRβ, NG2) via neural crest cells (NC, marker Sox10). B) Gene expression of pericyte markers (*PDGFRB, MCAM —* encoding CD146*, CSPG4* — encoding NG2) in *APOE3/E3* iPSCs, NC, and pericytes by qRT-PCR. C) Expression of NG2 and PDGFRβ in *APOE3/E3* pericytes. D) Gene expression of pericyte markers (*PDGFRB, MCAM —* encoding CD146*, CSPG4* — encoding NG2) in *APOE2/E2* iPSCs, NC, and pericytes by qRT-PCR. E) Expression of NG2 and PDGFRβ in *APOE2/E2* pericytes. F) Gene expression of pericyte markers (*PDGFRB, MCAM —* encoding CD146*, CSPG4* — encoding NG2) in *APOE4/E4* iPSCs, NC, and pericytes by qRT-PCR. G) Expression of NG2 and PDGFRβ in *APOE4/E4* pericytes. H) Gene expression of pericyte markers (*PDGFRB, MCAM —* encoding CD146*, CSPG4* — encoding NG2) in *APOE* KO iPSCs, NC, and pericytes by qRT-PCR. I) Expression of NG2 and PDGFRβ in *APOE* KO pericytes. Nuclei were counterstained with DAPI (blue). Scale bars 100 µm. Points in graphs represent individual replicates (cells from independent differentiations, n = 3-4), bars represent the mean, and error bars ± SD, *p < 0.05, **p < 0.01, ***p < 0.001, ****p < 0.0001, ns — not significant.

The expression of pericyte markers *PDGFRB* and *MCAM* (encoding CD146) (Fig. 1 B, D, F, H) in the resulting cells, together with the loss or downregulation of pluripotency markers *POU5F1*, *SOX2*, *NANOG* (Sup. Fig. 1 A, C, E, G), and the NC marker *SOX10* (Sup. Fig. 1 B, D, F, H) confirmed the differentiation towards pericytes. Notably, despite the increased expression of *PDGFRB* and *MCAM* in all *APOE-*isogenic pericytes, *CSPG4* (encoding NG2) expression was variable with no significant increase in *APOE3/E3* and *APOE4/E4* pericytes compared to iPSCs (Fig. 1 B, D, F, H). We then compared the expression of pericyte markers between all *APOE*-isogenic pericytes. Despite variability between genotypes, we did not observe a single genotype with significantly lower expression of all markers (Sup. Fig. 2 A). Similarly, all iPSC-derived *APOE*-isogenic pericytes expressed NG2 and PDGFRβ proteins, regardless of morphological variability (Fig. 1 C, E, G, I).

**Figure 2.**
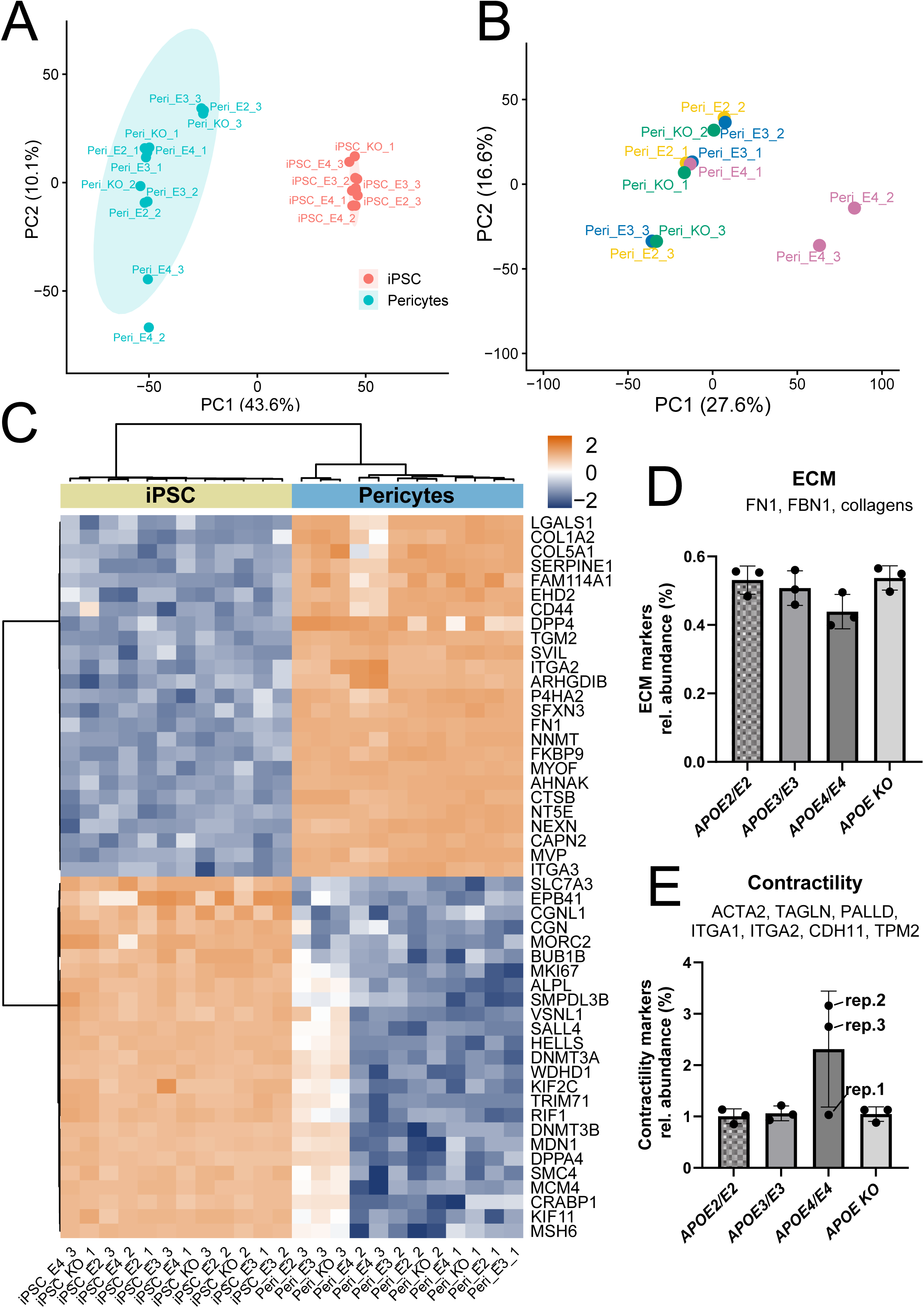
Characterization of iPSC and iPSC-derived pericyte proteotypes. A) Principal component analysis (PCA) of iPSC and pericyte protein profiles. Each point represents an individual biological replicate, colored by cell type. B) PCA of protein profiles of *APOE*-isogenic pericytes. Each point represents an individual biological replicate, colored by *APOE* genotype. PC1 and PC2 explain X% and Y% of the total variance, respectively. C) Heatmap represents color-coded abundance (Z-scored, row-scaled) of top 50 differentially abundant (FDR < 0.05, ranked by |log_2_FC|) proteins between iPSCs and iPSC-derived pericytes. D-E) Relative abundance of extracellular matrix (ECM) proteins (D) and contractility-linked proteins (E) in *APOE*-isogenic pericytes. Relative abundances were calculated as the percentage contribution of each protein signal to the total signal intensity of all proteins per sample. Points in graphs represent individual replicates (n = 3), bars represent the mean, and error bars ± SD.

To further investigate the gene expression variability between pericytes with different *APOE* genotypes, we characterized the proteotypes of *APOE*-isogenic iPSCs and iPSC-derived pericytes using LC-MS/MS proteomics. Principal component analysis (PCA) revealed cell type-specific clustering of the investigated cells, with more variability observed between iPSC-derived pericytes than between iPSCs. Among the iPSC-derived pericytes, one *APOE4/E4* replicate (Peri_E4_2) was separated from the main pericyte cluster along PC2, which accounted for only 10.1% of the total variance, however its position along PC1 remained within the range of the other pericytes (Fig. 2 A). To investigate this difference further, we compared all iPSC-derived *APOE*-isogenic pericytes and observed that PCA did not clearly cluster them by genotype. Two *APOE4/E4* replicates (Peri_E4_2 and Peri_E4_3) showed divergence, albeit with limited variance explained by PC1 (16.6%) and PC2 (27.6%) (Fig. 2 B). We then performed differential expression analysis to identify the proteins that are most characteristic of iPSCs and iPSC-derived pericytes. We found that proteins linked to the cell cycle and gene expression regulation, such as the proliferation marker protein Ki-67 and DNA (cytosine-5)-methyltransferase 3A, were upregulated in iPSCs. In contrast, pericytes expressed proteins related to cell adhesion and ECM organization, including fibronectin (FN1), galectin-1 (LGALS1), and nexilin (NEXN) (Fig. 2 C). Genes encoding these proteins have been previously identified in the brain mural cell transcriptome [17]. The levels of cell type-specific proteins were similar across different *APOE* genotypes and biological replicates in both iPSCs and pericytes. The Peri_E4_2 (APOE4/E4 pericytes replicate 2) and Peri_E4_3 (APOE4/E4 pericytes replicate 3) identified by PCA exhibited reduced expression of proteins associated with cell adhesion, including galectin-1 (LGALS1), collagens (COL1A2 and COL5A1), and CD44 [18]. Additionally, they had lower expression of the serine protease inhibitor SERPINE1, multifunctional regulatory protein FAM114A1, and ATP– and membrane-binding protein EHD2. Recent evidence suggests that *APOE4* pericytes have a myofibroblast-like phenotype, characterized by high expression of ECM proteins, such as FN1 and collagens, combined with elevated expression of proteins linked to cell contractility [6]. Based on this evidence, we calculated the relative abundances of ECM and contractility-linked proteins in *APOE-*isogenic pericytes. We observed a slight decrease in ECM proteins (Fig. 2 D) and an increase in the abundance of contractility-linked proteins in *APOE4/E4*, particularly in the two most divergent replicates (Fig. 2 E). However, statistical testing revealed no significant differences in ECM or contractility protein abundance between different pericyte genotypes. Overall, despite variability among individual pericyte replicates, all iPSC-derived pericytes shared a characteristic pericyte proteotype. They did not cluster by *APOE* genotype, indicating that the variability was not primarily driven by apoE.

Next, we confirmed that *APOE2/E2*, *APOE3/E3*, and *APOE4/E4* pericytes expressed *APOE*, while *APOE* KO cells did not (Sup. Fig. 2B). Furthermore, the *APOE*-expressing pericytes secreted apoE, with the highest levels observed in *APOE3/E3* pericyte-conditioned media. ApoE was not detectable in *APOE* KO pericyte-conditioned media and vehicle media that had no contact with the cells (Sup. Fig. 2 C).

### 3.2 Lipid composition of LpEs secreted by pericytes is specific to *APOE* **genotype**

Previous studies by our group and others have reported that the protein and lipid compositions of LpEs are cell type-specific [8,10]. To investigate whether pericyte-secreted LpE compositions also depend on the *APOE* genotype, we cultured *APOE*-isogenic pericytes in delipidated media for three days. We then collected the conditioned media and immunoprecipitated LpEs for lipid profiling. A total of 98 lipid species were identified. After filtering out the lipids enriched in vehicle media and in *APOE* KO pericyte-conditioned media, 46 lipids were identified as LpE components (Fig. 3 A). All particle lipids were highly abundant LpE4, while LpE2 and LpE3 were less lipidated and exhibited different lipidation patterns (Fig. 3 B). Differential expression analysis of lipids associated with LpE3 and LpE2 revealed that ether-linked phosphatidylcholines (PC-O) and phosphatidylethanolamines (PE-O) were more abundant in LpE3, whereas ester-linked phosphatidylcholines (PC) and sphingomyelins (SM) were more prevalent in LpE2. PC O-36:2, PE O-38:4, and PC O-38:5 were among the most significantly upregulated lipid species in LpE3, while short-chain PC 32:2, PC 32:1, and SM 34:1;O2 were the most significantly upregulated in LpE2 (Fig. 3 C). Lipid subclass enrichment analysis further confirmed the higher representation of the PC-O and PE-O subclasses in LpE3, and SM and PC subclasses — in LpE2 (Fig. 3 D). Comparisons of LpE4 with either LpE3 or LpE2 revealed that all identified lipid subclasses were enriched in LpE4 (Fig. 3 E-F). We then calculated the absolute concentrations of lipids in each condition and confirmed that the concentrations of all lipid subclasses were the highest in LpE4 (Sup. Fig. 3). Next, we calculated the relative abundance of each lipid subclass to investigate the compositions of pericyte-secreted LpEs independently of total lipid levels. We observed that the contributions of PC, PC-O, and SM to the total lipids were the highest in all genotypes. PC accounted for 81.3% of lipids detected in LpE2, 70.0% in LpE3, and 72.3% in LpE4. Of the three most abundant lipid subclasses, PC-O was the most divergent across genotypes. LpE2 contained markedly less PC-O (1.3%) than LpE3 and LpE4 (8.8% and 7.9%, respectively). In contrast, SM contribution was the highest in LpE2 with 11.6%, compared to 9.8% in LpE3 and 7.5% in LpE4. PE, PE-O, triglycerides (TG), phosphatidylserine (PS), and phosphatidylinositol (PI) were less abundant, each subclass accounting for less than 4% of the total lipids detected in LpE. However, their abundances differed between genotypes. PE and PE-O were less represented in LpE2 compared to other particles, whereas TG, PS, and PI were the most abundant in LpE4, followed by LpE3, and LpE2 (Fig. 3 G).

**Figure 3.**
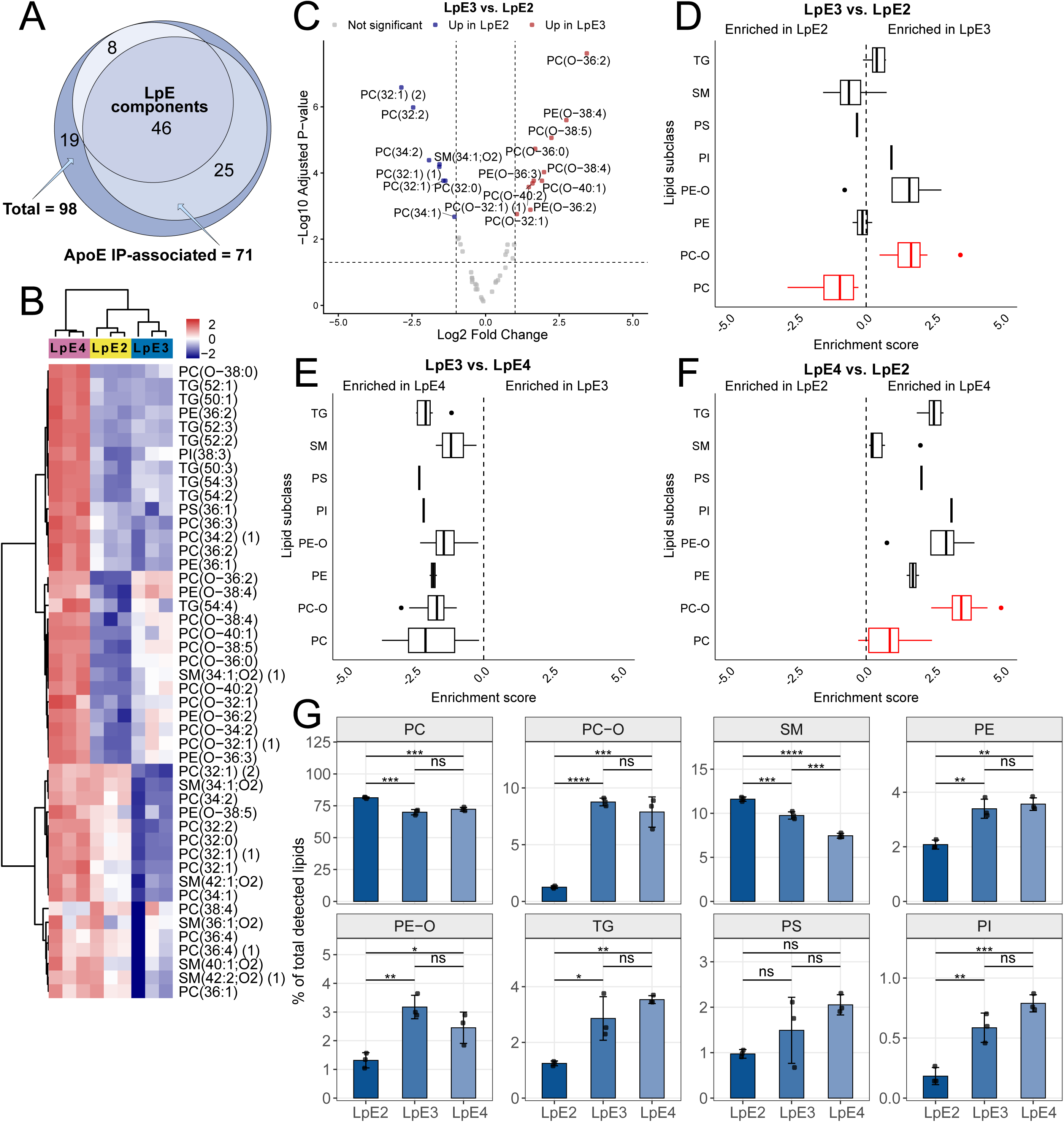
Lipid profiling reveals different lipidation patterns of pericyte-secreted LpEs depending on *APOE* genotype. A) The overlap of all lipids identified in the study, lipids identified as associated with apoE immunoprecipitation (ApoE IP-associated; Enriched (FC ≥ 1.5, FDR < 0.05) in each experimental condition vs. vehicle medium), and LpE components (LpE components; Enriched (FC ≥ 1.5, FDR < 0.05) in each experimental condition vs. *APOE* KO conditioned medium). B) Heatmap represents color-coded abundance (Z-scored, row-scaled) of lipids identified as components of pericyte-secreted LpE3, LpE2, and LpE4. C) Differential expression of lipids identified as components of pericyte-secreted LpE3 vs. LpE2. Each point represents one lipid species. Significance thresholds: |log_2_FC| ≥ 1 (FC ≥ 2), FDR < 0.05. D-F) Lipid subclass enrichment analysis in LpEs secreted by *APOE3/E3* vs. *APOE2/E2* (D), *APOE3/E3* vs. *APOE4/E4* (E), *APOE4/E4* vs. *APOE2/E2* (F) pericytes. Lipid subclass distributions are shown as boxplots with medians as center lines, whiskers extend to the extreme non-outlier values, dots represent outliers. Significantly different subclasses (FDR < 0.05) are highlighted. G) Relative abundances of lipid subclasses in LpEs secreted by *APOE2/E2* (LpE2), *APOE3/E3* (LpE3), and *APOE4/E4* (LpE4) pericytes. Relative abundances were calculated as the percentage contribution of each lipid subclass signal to the total signal intensity of all lipids per sample. Only lipids identified as LpE components were used in the calculations. Points in graphs represent individual replicates (n = 3), bars represent the mean, and error bars ± SD. Pairwise comparisons were performed within each lipid subclass, *p < 0.05, **p < 0.01, ***p < 0.001, ****p < 0.0001 (adjusted p-values), ns — not significant.

After identifying the lipid components of pericyte-secreted LpE, we aimed to verify their presence in human lipoproteins *in vivo.* To do so, we referred to the CSF-derived LpE composition data from our previous study [10]. Twenty-three of the 40 lipids that we identified as pericyte derived LpE components (isobaric species were treated as a single lipid) were previously noted in our CSF-derived LpE lipid profiling dataset (Sup. Table 1). The remaining lipid species were individually low in abundance, accounting for less than 2% of the total lipids detected in particles of any genotype. Together, they comprised 6.3% of lipids detected in LpE2, 11.9% of LpE3, and 11.4% of LpE4 lipids (Sup. File 3 – relative abundances of LpE lipids).

### 3.3 Pericyte cellular lipids vary depending on *APOE* genotype

To investigate whether genotype-dependent differences in the lipid composition of pericyte-secreted LpEs reflect differences in cellular lipid composition, we performed lipid profiling of *APOE*-isogenic pericytes. Lipids defined as LpE components were most abundant in *APOE3/E3* cells, followed by *APOE* KO pericytes (Sup. Fig. 4 A). We then ranked the lipid species of LpEs and cells for comparison of their relative abundances (Sup. Fig. 4 B and C) (Sup. File 3 – relative abundances of LpE lipids, Sup. File 4 – relative abundances of cellular lipids). Six out of the top 10 ranked lipids in both lists overlapped, namely PC 34:1, PC 32:0, PC 36:1, PC 36:2, SM 34:1;O2, and PC 32:1. We then calculated the absolute concentrations of these lipids in both pericyte-secreted LpEs (Sup. Fig. 4 D) and cells (Sup. Fig. 4 E). All six lipids were the most abundant in LpE4 and in *APOE3/E3* pericytes.

**Figure 4.**
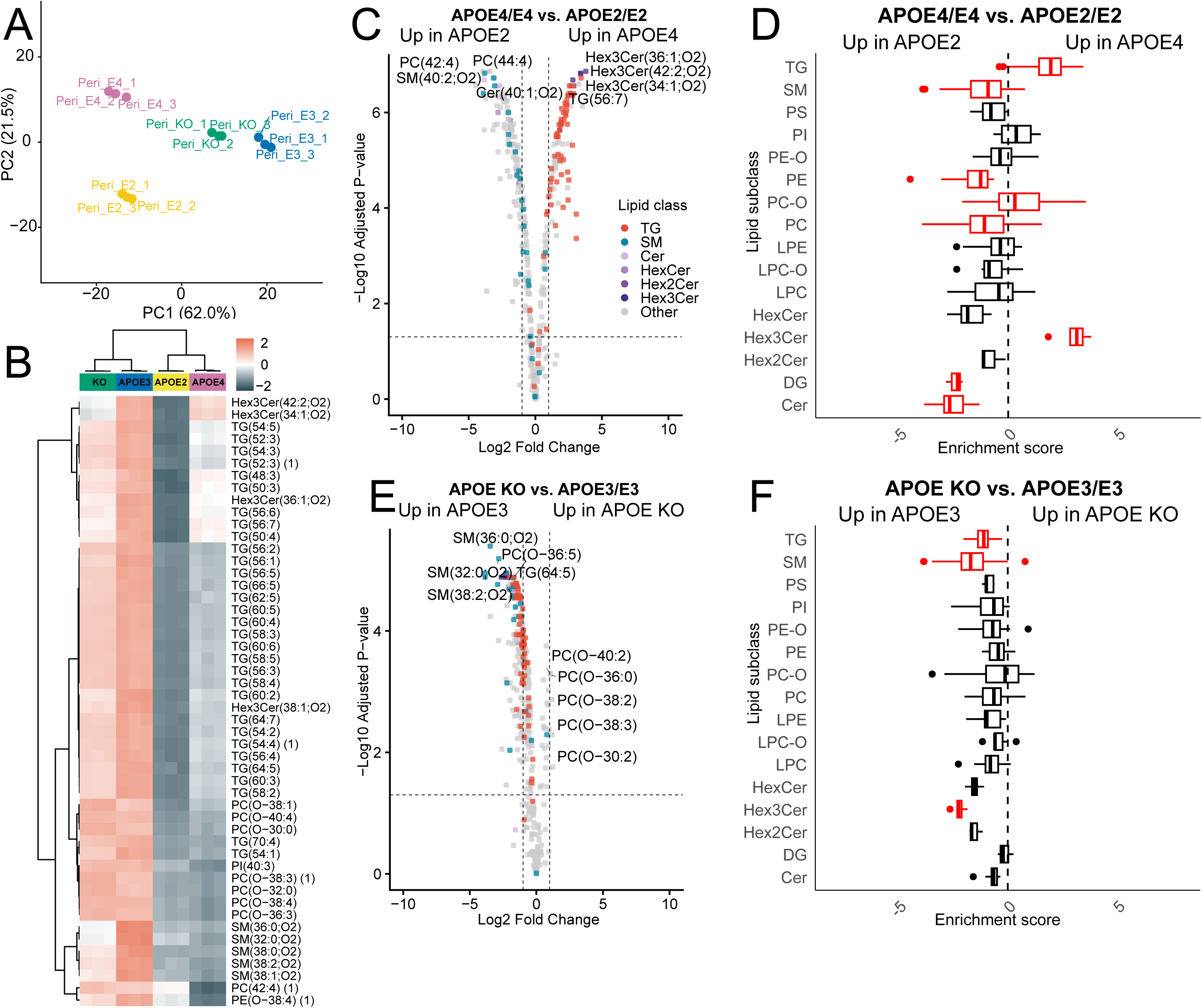
Cellular lipid profiling reveals different lipidation patterns of pericytes depending on *APOE* genotype. A) PCA of lipid profiles in *APOE*-isogenic pericytes. Each point represents an individual biological replicate, colored by *APOE* genotype. PC1 and PC2 explain X% and Y% of the total variance, respectively. B) Heatmap represents color-coded abundance (Z-scored, row-scaled) of cellular lipids in *APOE3/E3*, *APOE2/E2*, *APOE4/E4*, and *APOE* KO pericytes. Top 50 lipid species with the largest effect size among lipids showing significant differential abundance across four groups are shown, ranked by |log_2_FC|. An omnibus moderated F-test was used to identify lipids differing significantly across any contrast (FDR < 0.05). C) Differential expression of cellular lipids in *APOE4/E4* vs. *APOE2/E2* pericytes. D) Lipid subclass enrichment analysis of cellular lipids in *APOE4/E4* vs. *APOE2/E2* pericytes. E) Differential expression of cellular lipids in *APOE* KO vs. *APOE3/E3* pericytes. F) Lipid subclass enrichment analysis of cellular lipids in *APOE* KO vs. *APOE3/E3* pericytes. Lipid subclass distributions are shown as boxplots with medians as center lines, whiskers extend to extreme non-outlier values, dots represent outliers. Significantly different subclasses (FDR < 0.05) are highlighted. Each point in differential expression plots represents one lipid species. Significance thresholds for differential expression plots: |log2FC| ≥ 1 (FC ≥ 2), FDR < 0.05. FC – fold change, FDR – false discovery rate.

We then proceeded to further investigate the cellular lipid profiles of *APOE*-isogenic pericytes. In contrast to the protein profiles, PCA of cellular lipids showed genotype-dependent clustering (Fig. 4 A). Generally, lipids were more abundant in *APOE3/E3* and *APOE* KO pericytes compared to *APOE2/E2* and *APOE4/E4* cells (Fig. 4 B). We identified TG as the lipid subclass most broadly affected by *APOE* genotype. Many TG species were enriched in *APOE3/E3* and *APOE* KO pericytes compared to *APOE2/E2* and *APOE4/E4* pericytes. However, several species such as TG 48:3, TG 50:3, TG 56:6, TG 56:7, and TG 50:4 were also abundant in *APOE4/E4* pericytes. Additionally, we observed genotype-dependent variation in the abundance of SM and trihexosylceramides (Hex3Cer) (Fig. 4 B). Next, we calculated the absolute concentration of each lipid subclass in pericytes of each genotype. The concentrations of almost all lipid subclasses were the highest in *APOE3/E3* pericytes, except for PE, which was the most abundant in *APOE2/E2* cells (Sup. Fig. 5).

**Figure 5.**
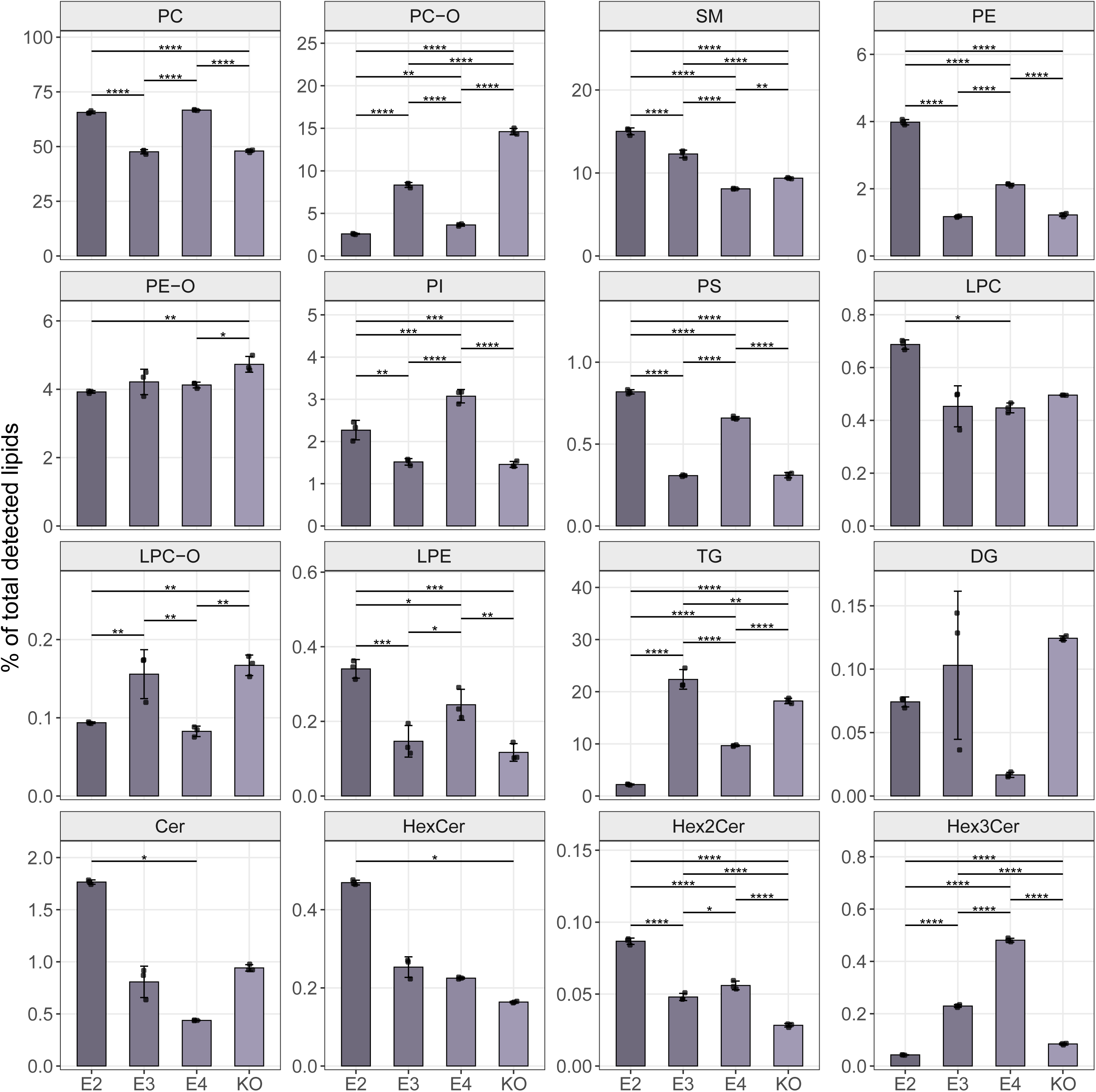
Relative abundances of cellular lipids in *APOE*-isogenic pericytes. Relative abundances of cellular lipid subclasses in *APOE2/E2* (E2), *APOE3/E3* (E3), and *APOE4/E4* (E4) pericytes. Relative abundances were calculated as the percentage contribution of each lipid subclass signal to the total signal intensity of all lipids per sample. Points in graphs represent individual replicates (n = 3), bars represent the mean, and error bars ± SD. Pairwise comparisons were performed within each lipid subclass, *p < 0.05, **p < 0.01, ***p < 0.001, ****p < 0.0001 (adjusted p-values). Only significant pairwise comparisons are shown.

Since we observed generally lower lipid levels in *APOE4/E4* and *APOE2/E2* pericytes compared to *APOE3/E3* and *APOE* KO, we first performed differential expression analysis of lipids in *APOE4/E4* versus *APOE2/E2* pericytes. We confirmed the upregulation of TG in *APOE4/E4*. Conversely, most SM species were upregulated in *APOE2/E2*. Several Hex3Cer species (Hex3Cer 42:2;O2, Hex3Cer 34:1;O2, Hex3Cer 36:1;O2) were among the most significantly upregulated lipids in *APOE4/E4*, whereas dihexosylceramides (Hex2Cer), hexosylceramides (HexCer), and ceramides (Cer), especially Cer 40:1;O2, were enriched in *APOE2/E2* (Fig. 4 C). These differences were confirmed by lipid subclass enrichment analysis in *APOE4/E4* versus *APOE2/E2* pericytes. Additionally, we observed higher levels of PE, PC, and diglycerides (DG) in *APOE2/E2*, and higher levels of PC-O in *APOE4/E4* (Fig. 4 D). Next, we performed differential expression analysis of lipids in *APOE* KO versus *APOE3/E3* pericytes and noted a higher representation of TG together with the enrichment of SM (especially SM 36:0;O2, SM 32:0;O2, and SM 38:2;O2) in *APOE3/E3*. Several PC-O species (PC O-40:2, PC O-36:0, PC O-38:2, PC O-38:3, and PC O-30:2) were upregulated in *APOE* KO pericytes (Fig. 4 E). However, lipid subclass enrichment analysis in *APOE* KO versus *APOE3/E3* cells showed enrichment of all lipid subclasses in *APOE3/E3*, including a slight shift in PC-O representation towards *APOE3/E3* (Fig. 4 F).

Next, we performed differential expression analysis of lipids in *APOE4/E4* pericytes in comparison with those of *APOE* KO and *APOE3/E3* cells, respectively. We observed a higher representation of most lipid species in *APOE* KO and *APOE3/E3* cells (Sup. Fig. 6 A, B). Several PC-O species (PC O-38:3, PC O-38:2, PC O-40:4, PC O-38:3, and PC O-40:5) were enriched in *APOE* KO cells, whereas the highly unsaturated PC O-30:5, PC O-46:6, and PC O-46:5 were upregulated in *APOE4/E4*. Hex3Cer 42:1;O2 and PI 38:6 were also among the most strongly upregulated lipids in *APOE4/E4* cells (Sup Fig. 6 A). PI 38:6 was also upregulated in *APOE4/E4* cells in comparison to *APOE3/E3* cells (Sup Fig 6 B). We then performed differential expression analysis of lipids in *APOE2/E2* pericytes in comparison with those in *APOE* KO and *APOE3/E3* cells, respectively. Similarly to the previous analyses including *APOE4/E4* cells, most lipid species were enriched in *APOE* KO and *APOE3/E3* pericytes. However, short-chain PE 30:1, long-chain LPC 24:0, highly unsaturated PC 40:6 and PE 36:4, short and unsaturated PC O-30:5 and PC O-30:4, and HexCer 42:1;O2 were upregulated in *APOE2/E2* (Sup. Fig. 6 C, D).

**Figure 6.**
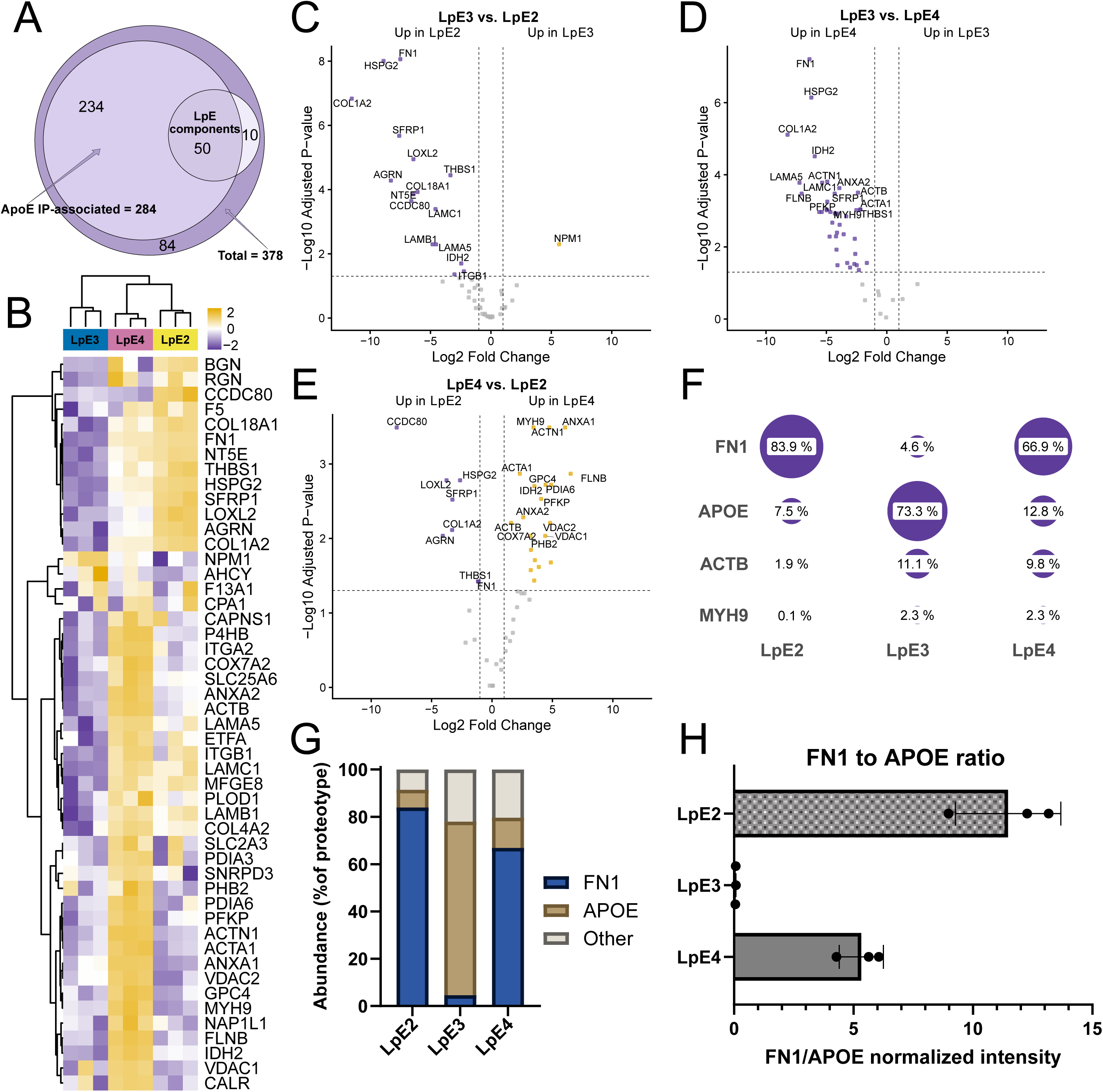
Proteotype profiling reveals different protein composition of pericyte-secreted LpEs depending on *APOE* genotype. A) The overlap of all proteins identified in the study, proteins identified as associated with apoE immunoprecipitation (ApoE IP-associated; Enriched (FC ≥ 2, FDR < 0.05) in each experimental condition vs. vehicle medium), and LpE components (Enriched (FC ≥ 2, FDR < 0.05) in each experimental condition vs. *APOE* KO conditioned medium). B) Heatmap represents color-coded abundance (Z-scored, row-scaled) of proteins in pericyte-secreted LpE3, LpE2, and LpE4. C-E) Differential expression of proteins identified as components of LpE3 vs. LpE2 (C), LpE3 vs. LpE4 (D), and LpE4 vs. LpE2 (E) secreted by pericytes. Each point represents one protein. Data are normalized to *APOE* intensity in addition to global (median-centered) normalization. Significance thresholds: |log2FC| ≥ 1 (FC ≥ 2), FDR < 0.05. F) Dot plot shows the abundances of proteins identified as LpE components across *APOE* genotypes. Protein abundances are expressed as the percentage contribution of each protein to the total protein signal. G) Protein composition of pericyte-secreted LpEs across *APOE* genotypes. Relative abundances calculated as the percentage contribution of *APOE*, FN1, and all other protein components to total particle proteotype. Abundances represented as mean of three replicates per *APOE* genotype. H) FN1 to APOE ratio in pericyte-secreted LpE, calculated as FN1/APOE normalized intensity, across *APOE* genotypes. Points in graphs represent individual replicates (n = 3), bars represent the mean, and error bars ± SD.

Next, we calculated the relative abundance of each lipid subclass to investigate the lipid profiles of *APOE*-isogenic pericytes independently of total lipid levels. The relative abundances of ceramides and glycosphingolipids confirmed a pattern previously observed in the *APOE4/E4* versus *APOE2/E2* differential expression analysis: Hex3Cer were the most abundant in *APOE4/E4* pericytes, while the relative abundances of Hex2Cer, HexCer, and Cer were the highest in *APOE2/E2*. TG were the most represented in *APOE3/E3*, followed by *APOE* KO, *APOE4/E4*, and *APOE2/E2* pericytes. Several lipid subclasses had elevated contributions in both *APOE3/E3* and *APOE* KO pericytes compared to *APOE2/E2* and *APOE4/E4* cells: PC-O, LPC-O, and DG. Conversely, PC, PI, PS, and LPE contributions were higher in *APOE2/E2* and *APOE4/E4* pericytes and lower in *APOE3/E3* and *APOE* KO. SM, LPC, and PE-O did not follow that pattern. SM abundance increased in the order of *APOE4/E4*<*APOE* KO<*APOE3/E3*<*APOE2/E2*. LPC contribution was the highest in *APOE2/E2* cells and broadly comparable across other genotypes. PE-O contribution was the highest in *APOE* KO and comparable across other genotypes (Fig. 5).

### 3.4 Protein composition of LpEs secreted by pericytes is specific to *APOE* **genotype**

Next, we isolated pericyte-secreted LpEs as described above and investigated their protein compositions using LC-MS/MS proteomics. A total of 378 proteins were identified in the analysis. After filtering out the proteins enriched in vehicle media and in the *APOE* KO pericyte-conditioned media, 50 proteins were identified as components of LpEs (Fig. 6 A). Most of these proteins were highly represented in LpE4, followed by LpE2. In contrast, LpE3 showed the least abundance of associated cargo proteins (Fig. 6 B). Differential expression analysis of proteins in LpE3 versus LpE2 revealed nucleolar protein nucleophosmin (NPM1) as significantly upregulated in LpE3 while several ECM proteins were significantly enriched in LpE2. Among the latter, we identified fibronectin-1 (FN1), perlecan (HSPG2), thrombospondin-1 (THBS1), collagen COL18A1, and laminin subunits (LAMC1, LAMB1, and LAMA5). FN1 and HSPG2 were the strongest differential features in this comparison (Fig. 6 C). LpE3 versus LpE4 comparison revealed that, similarly to LpE2, LpE4 are most strongly enriched with FN1 and HSPG2, while no proteins were upregulated in LpE3. (Fig. 6 D). Differential expression analysis of LpE4 versus LpE2 components revealed the upregulation of coiled-coil domain-containing protein 80 (CCDC80), which promotes cell adhesion and matrix assembly, together with agrin (AGRN), THBS1, FN1, and HSPG2 in LpE4. STRING Gene Ontology (GO) search pointed to regulation of cell-substrate adhesion and angiogenesis as terms associated with these proteins [18]. Among features upregulated in LpE4, we noted annexins (e.g., ANXA1, ANXA2), actins (e.g., ACTB, ACTA1), and myosin-9 (MYH9) (Fig. 6 E). STRING GO search revealed actin cytoskeleton organization as the main term associated with them [18]. We then compared the protein components of pericyte-secreted LpEs with those in human lipoproteins isolated from CSF that were previously reported by three studies [9,19], including one by our group [10]. Of the 50 proteins recovered in pericyte-derived LpE, three were identified in CSF-derived LpEs by all studies: APOE, FN1, and coagulation factor V (F5). In addition, lactadherin (MFGE8), adenosylhomocysteinase (AHCY), LAMC1, MYH9, secreted frizzled-related protein 1 (SFRP1), and THBS1 were noted in our previous investigation of CSF-derived LpEs [10]. ANXA2, HSPG2, collagen COL1A2, ACTB, and calreticulin (CALR) were also reported by one or two of the studies by other groups[9,19]. The other 36 proteins in pericyte-derived LpEs were not observed in human CSF-derived LpEs by the studies included in this comparison (Sup. Table 2).

Using relative abundance calculations, we observed that FN1 and APOE were the dominant components of pericyte-derived LpEs (Fig. 6 F). However, their relative abundance varied by apoE isoform: FN1, APOE, and other proteins accounted for 83.9%, 7.5%, and 8.6% of LpE2, respectively; 66.9%, 12.8%, and 20.3% of LpE4; 4.6%, 73.3%, and 22.1% of LpE3. (Fig. 6 G). Notably, by normalizing the FN1 intensity signal to APOE, we confirmed that the genotype-dependent differences in FN1 abundance were independent of the amount of apoE recovered by immunoprecipitation and detected by LC-MS/MS (Fig. 6 H). ACTB was among the most abundant proteins in LpE3 (11.1%) and LpE4 (9.8%), but not LpE2 (1.9%) (Sup. File 5 – relative abundances of particle proteins).

Because FN1 levels differed markedly in LpEs depending on the genotype, we next investigated cellular FN1 levels in *APOE*-isogenic pericytes. FN1 was among the top 100 most abundant proteins in all investigated conditions. However, its cellular abundance did not differ substantially across *APOE* genotypes: FN1 comprised 0.320% of the *APOE2/E2* pericyte proteotype (ranked 59^th^ of 5127 features by relative abundance), 0.326% of *APOE3/E3* (54^th^), 0.285% of *APOE4/E4* (65^th^), and 0.320% of the *APOE* KO proteotype (58^th^) (Sup. Fig. 7 A) (Sup. File 6 – relative abundances of cellular proteins). Observed variability in FN1 expression between replicates was not genotype-specific (Sup. Fig. 7 B).

## 4. Discussion

In this study, we demonstrate that the protein and lipid components of LpEs secreted by brain pericytes vary depending on the *APOE* genotype. The distinct genotype-dependent cargo of pericyte-secreted LpEs points to potentially different roles of these particles in brain homeostasis and neurodegenerative diseases. Additionally, the abundance and composition of cellular lipids are influenced by the *APOE* genotype (or the absence of *APOE*), while differences in the proteomes of *APOE*-isogenic pericytes are more modest. Notably, genotype-dependent patterns in the levels and composition of lipids in pericyte-secreted LpEs do not mirror those observed in cellular lipids, suggesting that the role of *APOE* in pericyte lipid metabolism extends beyond lipid transport via LpEs.

Pericyte-derived LpE4 contained more of the investigated lipids than LpE2 and LpE3. This finding was unexpected given the conformation of the lipid-binding domain in apoE4 is thought to have a reduced lipid-binding capacity [20]. Previous studies using native-PAGE have described LpE4 as less lipidated than LpE3 in human iPSC-derived astrocytes [21] and in mice overexpressing the human apoE4 in astrocytes [22]. Conversely, apoE4 has been identified as the most lipidated isoform secreted by human immortalized astrocytes and embryonic kidney cells [23]. Another study has suggested that, rather than having decreased lipid content, LpE4 contain more apoE molecules per particle than LpE3 [24]. Additionally, apoE4 preferentially binds to larger lipoproteins, while apoE3 binds to smaller ones, which could also result in higher levels of apoE4-bound lipids [25]. Finally, we focused on sphingolipids, glycerophospholipids, and triglycerides, which can be reliably profiled together using a single LC-MS/MS method. Given that cholesterol is also a crucial component of brain-derived lipoproteins [7], future studies investigating cholesterol and cholesteryl esters in pericyte-secreted LpEs would complement these findings.

We noted minimal relative differences in lipid subclass levels between pericyte-secreted LpE3 and LpE4. It is consistent with recent findings by Strickland *et al.* [8] stating that cellular origin affects LpE composition more than the apoE isoform. However, LpE2 had reduced levels of PC-O and increased levels of SM. SM contribute to the organization and rigidity of the lipoprotein surface [26] and interact with cholesterol [27]. The increase of SM in LpE2 may therefore influence cholesterol organization, accessibility, and exchange. Relative TG levels in LpEs increased in the order of APOE2 < APOE3 < APOE4. Since triglyceride-rich lipoproteins and the products of their lipolysis (e.g., triglycerides and free fatty acids) increase BBB permeability [28], lower TG levels could reflect the protective role of pericyte-secreted LpE2.

In contrast to the strong lipidation of LpE4, we observed an overall increase of cellular lipids in *APOE3/E3* pericytes compared to both *APOE2/E2* and *APOE4/E4* cells. This indicates that the pronounced lipidation of LpE4 is not simply due to higher cellular lipid availability or facilitated lipid transfer to apoE. Notably, total lipid levels were similar in *APOE3/E3* and *APOE* KO cells, suggesting that *APOE2* and *APOE4* may affect pericyte lipid metabolism more than the complete absence of apoE.

TG, SM, and glycosphingolipids (Hex3Cer, Hex2Cer, HexCer) were among the most differentially abundant lipid subclasses between pericytes of different *APOE* genotypes. These lipid subclasses were also recently identified as the most divergent in the plasma lipidomes of *APOE3* and *APOE4* knock-in mouse models [29]. Previous studies reported increased accumulation of TG-enriched lipid droplets in *APOE4* astrocytes and microglia and linked it to higher inflammation and disease phenotypes [30,31]. In contrast, we did not observe a significant increase in TG in *APOE4/E4* pericytes. Together this demonstrates a fundamental difference in the effect of *APOE4* on lipid metabolism between cell types. Although TG levels were the highest in *APOE3/E3* pericytes, combined with high levels of other lipid subclasses this could reflect greater overall lipidation rather than the specific accumulation of the lipid droplet-associated TG in *APOE3/E3*. A recent study reported elevated TG, together with altered expression of proteins linked to lipid transport and degradation in pericytes carrying the *APOE2* allele, which is associated with reduced AD risk [32]. We did not observe similar patterns in our model. However, the authors of this study characterized iPSC-derived pericytes differentiated via the mesoderm, not the neural crest route. Therefore, this discrepancy may reflect the differences in pericyte origin and the heterogeneity of CNS pericytes.

Hex3Cer, although also the most abundant in *APOE3/E3* cells, were the most represented in *APOE4/E4* pericytes relative to total lipid content. Impaired glycosphingolipid processing, including reduced Hex3Cer catabolism has been linked to neurodegeneration, and particularly to Parkinson’s disease [33,34]. Glycosphingolipids are also associated with oxidative stress, mitochondrial dysfunction, and inflammation, which frequently coexist with neurodegeneration [35]. The downstream products of Hex3Cer catabolism (Hex2Cer, HexCer, and Cer) were more abundant in *APOE2/E2* pericytes relative to the total lipid content. This implies differences in glycosphingolipid metabolism between *APOE4/E4* and *APOE2/E2*, with potentially more efficient catabolism in *APOE2/E2*.

Notably, PI 38:6, one of the most upregulated lipids in *APOE4/E4* cells in comparison to other genotypes, has also been associated with oxidative stress responses [36]. Together with the enrichment of Hex3Cer, this may indicate increased oxidative stress in *APOE4/E4* pericytes, consistent with previous evidence linking apoE4 to increased amyloid-associated oxidative stress [37] and reduced protection against oxidative species compared to apoE3 [38]. Thus, although *APOE4/E4* pericytes do not show the lipid droplet-associated TG accumulation reported in *APOE4*-carrying astrocytes and microglia, their lipid profile points to potentially altered oxidative stress responses.

ECM proteins were the most abundant protein group in pericyte-derived LpEs. The abundance of these proteins was genotype-dependent, with the highest levels in LpE4. ECM proteins have been proposed as components of amyloid deposits, alongside amyloid fibrils and apoE [39]. They also contribute to the accumulation and stabilization of amyloid-β (Aβ) within the brain vasculature [40]. Cerebral amyloid angiopathy (CAA), characterized by the deposition of Aβ in the walls of cerebral blood vessels, is particularly prevalent in *APOE4* carriers [41]. *APOE2*, despite its protective effect against AD, has been associated with an increased risk of CAA-related hemorrhagic complications and vasculopathic changes in the affected vessels [42,43]. Thus, the preferential enrichment of ECM proteins in LpE4 and LpE2 may be relevant to their distinct effects on vascular amyloid pathology.

Among the ECM proteins, FN1 was of particular interest due to its high abundance in pericyte-secreted LpE2 and LpE4, but not in LpE3. While cellular FN1 levels were similar across *APOE*-isogenic pericytes, FN1 incorporation was notably higher in LpE4 and LpE2, suggesting that *APOE* genotype may influence lipoprotein assembly and composition. Excessive FN1 secretion by *APOE4/E4* vascular cells has been associated with increased amyloid deposition [6]. Furthermore, rare loss-of-function variants in FN1 have been linked to improved cognitive outcomes in *APOE4* carriers [44], supporting the notion of a functional interaction between FN1 and apoE in disease processes. Lastly, a recent study identified excessive FN1 accumulation as a key factor leading to *APOE4*-driven BBB leakage in AD. The study also noted the colocalization of FN1 and apoE4 [45]. Although the mechanisms underlying these associations remain unknown, increased FN1 content in pericyte-derived LpE4 and LpE2 suggests that FN1-containing LpEs may contribute to genotype-dependent differences in vascular amyloid deposition and BBB dysfunction.

There are a few methodological limitations of this study. First, our immunoprecipitation approach could not distinguish between lipidated, lipid-free, and extracellular vesicle-associated apoE. We considered the possibility that high levels of lipids observed in LpEs might be due to co-immunoprecipitation of apoE4 associated with lipid-rich vesicles. However, cellular lipid profiling did not show elevated lipidation of *APOE4/E4* cells. We also considered the possibility that differences in antibody recognition, such as isoform-dependent variation in antibody affinity during immunoprecipitation or apoE detection by ELISA, could explain the high lipidation of LpE4. However, our proteotype analysis revealed similar levels of apoE in different samples, suggesting that the observed differences are not due to technical limitations. Second, the *APOE4/E4* pericyte proteotype differed from its isogenic counterparts, although it was not consistent across all replicates. The differences included increased abundance of proteins linked to cell contractility. This finding is consistent with a recent study that reported an upregulation of contractility genes during the transition of *APOE4*-carrying cells toward a myofibroblast-like phenotype, which contributes to cerebrovascular pathology [6]. However, unlike the authors of this study, who also reported upregulated ECM genes, we observed a slight decrease in typical ECM protein abundance in *APOE4/E4* pericytes. Therefore, our findings partially support the *APOE4/E4* shift toward a myofibroblast-like state.

In summary, our findings demonstrate that pericyte-secreted LpEs differ in their lipid and protein composition depending on the *APOE* genotype. Therefore, they may perform distinct functions in brain homeostasis and in neurodegenerative disease pathology. Additionally, the lipidation and lipid subclass composition of cellular lipids in pericytes also varied in a genotype-dependent manner. This suggests that *APOE-*mediated lipid metabolism in vascular cells plays a role in brain health and disease that goes beyond the transport function of secreted LpEs.

## 5. Acknowledgements

JR is supported by a grant from the Synapsis Foundation for Dementia Research (No. 2022-PI04), a grant from the Bright Focus Foundation (A2021037S), and a grant from the Jubilee Foundation of the Swiss Life Insurance and Pension Institute for Public Health and Medical Research (Swiss Life Jubiläumsstiftung). Illustrations were made with templates from NIH BioArt Source (https://bioart.niaid.nih.gov/), Bioicons (https://bioicons.com/), and BioRender.com (License Robert_2026). We thank the team at the Functional Genomics Center for their support with lipidomics.

## 7. Data availability

The mass spectrometry proteomics data generated in this study have been submitted to the ProteomeXchange Consortium via its partner repository MassIVE, with the dataset identifier MSV000103573 and ProteomeXchange identifier PXD085591. The mass spectrometry lipidomics data have been submitted to MassIVE, with the data identifier MSV000103638.

## 8. Author contributions

Conceptualization — JR and EV; Data collection — EV, SG, AJ, SHA, SH; Data curation and analysis — EV, SG, AJ, AO; Methodology — EV, SG, AJ, Validation and visualization — EV; Supervision and funding acquisition — JR and AvE; Writing of the first draft — EV; Review and editing — all authors.

## 9. Figure captions

**Supplementary Figure 1.**
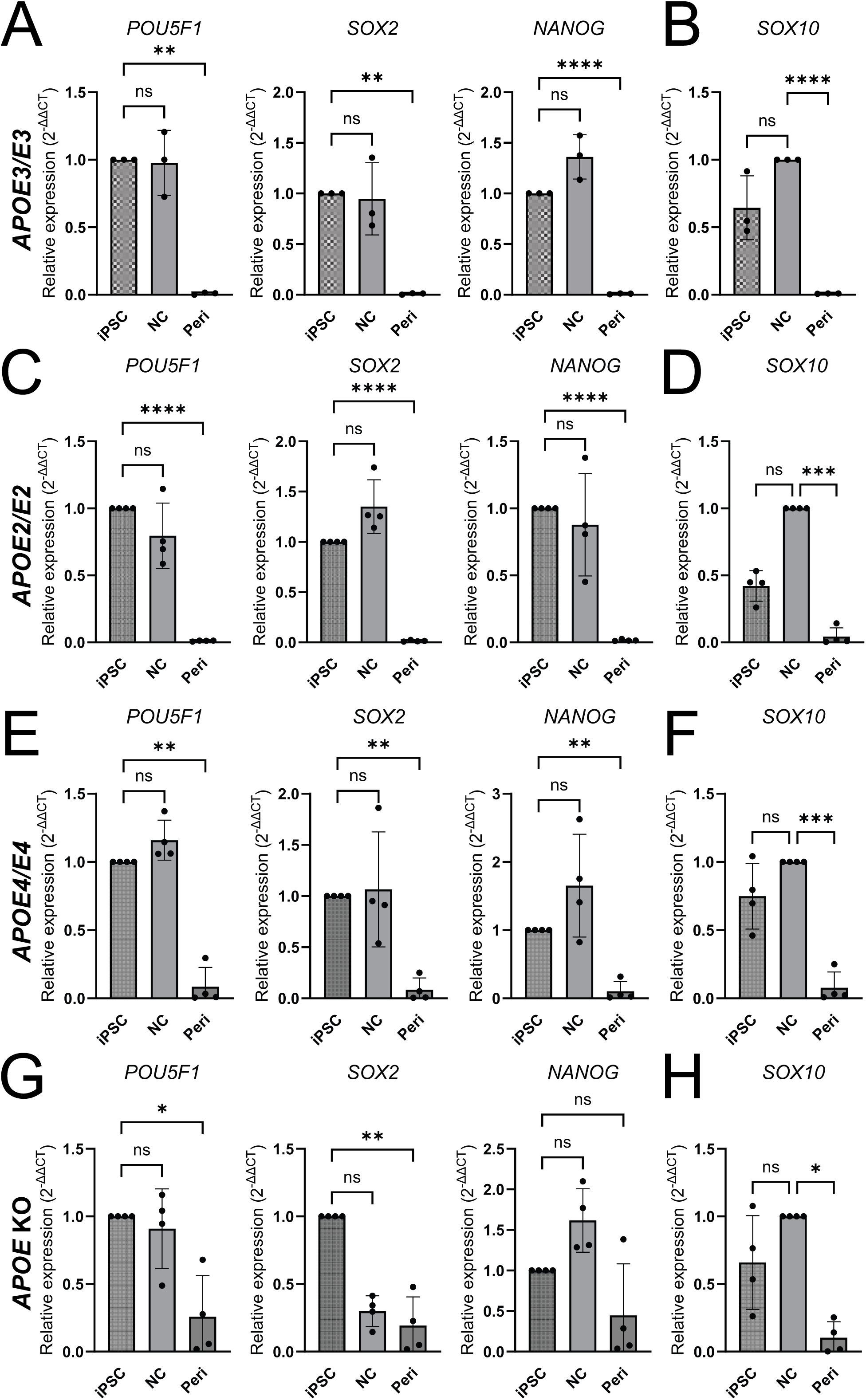
Relative gene expression of pluripotency and neural crest markers in iPSCs, neural crest cells, and iPSC-derived pericytes. A) Gene expression of pluripotency markers (*POU5F1*, *SOX2*, *NANOG*) and (B) the neural crest marker (*SOX10*) in *APOE3/E3* iPSCs, NC, and pericytes. C) Gene expression of pluripotency markers (*POU5F1*, *SOX2*, *NANOG*) and (D) the neural crest marker (*SOX10*) in *APOE2/E2* iPSCs, NC, and pericytes. E) Gene expression of pluripotency markers (*POU5F1*, *SOX2*, *NANOG*) and (F) the neural crest marker (*SOX10*) in *APOE4/E4* iPSCs, NC, and pericytes. G) Gene expression of pluripotency markers (*POU5F1*, *SOX2*, *NANOG*) and (H) the neural crest marker (*SOX10*) in *APOE* KO iPSCs, NC, and pericytes. Points in graphs represent individual replicates (n = 3-4), bars represent the mean, and error bars ± SD, *p < 0.05, **p < 0.01, ***p < 0.001, ****p < 0.0001, ns — not significant.

**Supplementary Figure 2.**
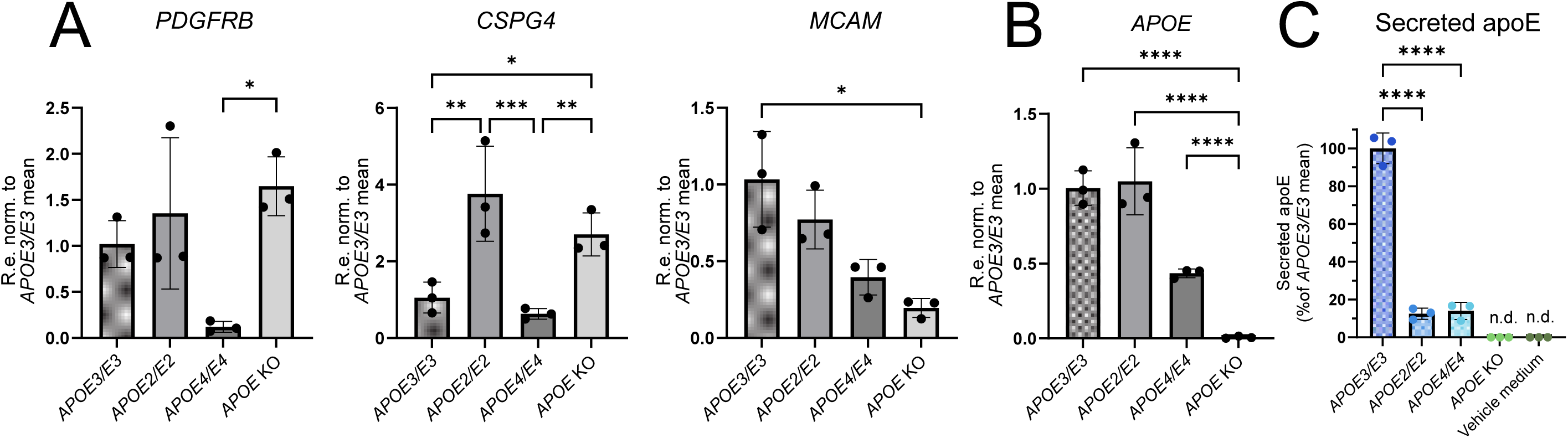
Relative gene expression of pericyte markers and *APOE*; ApoE secretion across iPSC-derived *APOE*-isogenic pericytes. A) Relative expression of pericyte markers (*PDGFRB, CSPG4* — encoding NG2, *MCAM —* encoding CD146) in *APOE-*isogenic pericytes, normalized to *APOE3/E3*. B) Relative expression of *APOE* in *APOE-*isogenic pericytes, normalized to *APOE3/E3*. Points in graphs represent individual replicates (n = 3), bars represent the mean, and error bars ± SD, *p < 0.05, **p < 0.01, ***p < 0.001, ****p < 0.0001. Only significant pairwise comparisons are shown. C) ApoE levels in conditioned media of *APOE*-isogenic pericytes, measured by ELISA, normalized to *APOE3/E3*. Points in graphs represent individual replicates (n = 3), bars represent the mean, and error bars ± SD, *p < 0.05, **p < 0.01, ***p < 0.001, ****p < 0.0001, n.d. — not detected (below assay detection limit). Only significant pairwise comparisons are shown.

**Supplementary Figure 3.**
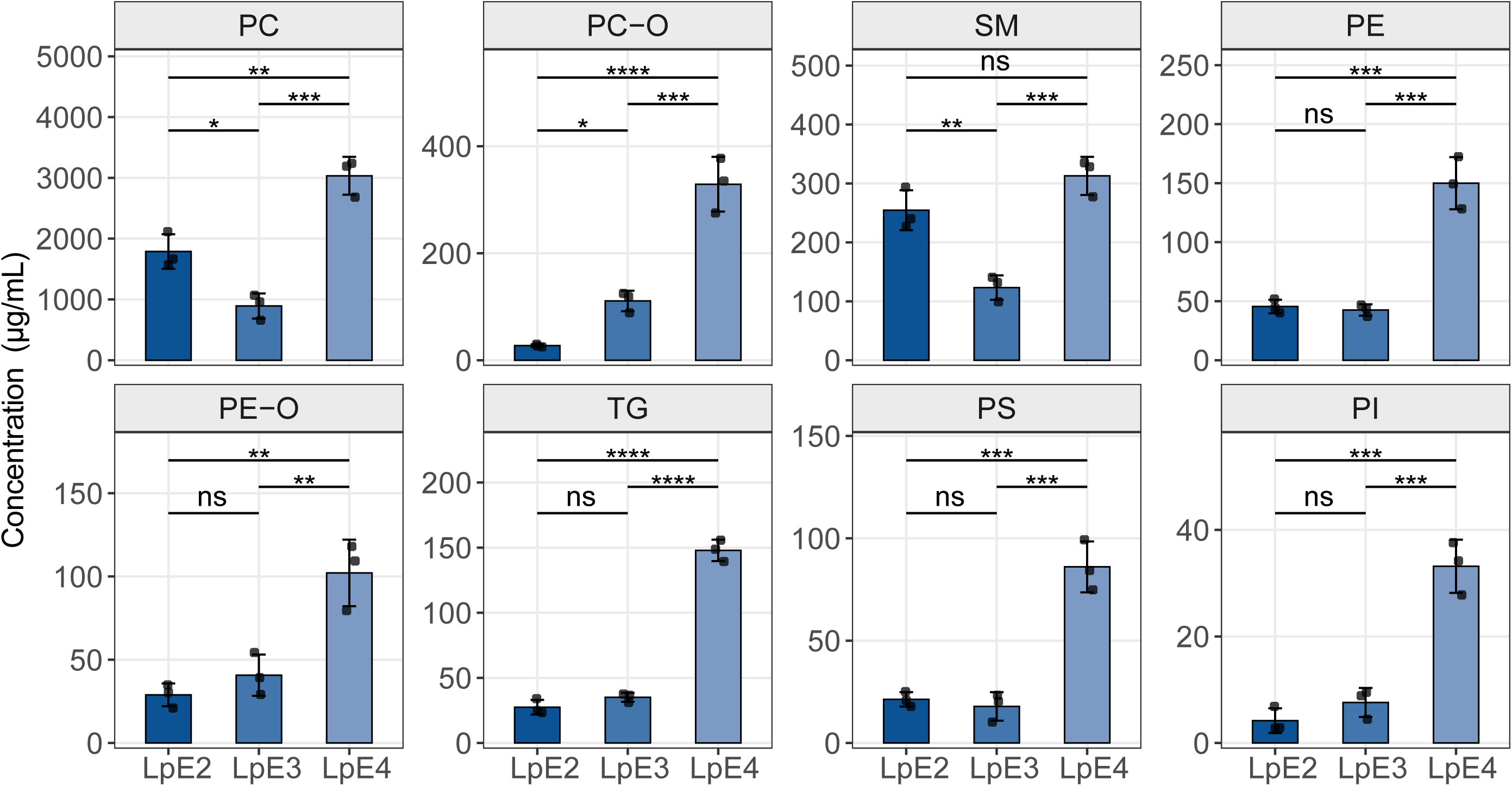
Lipid subclass concentrations in LpE. Absolute concentrations of lipid subclasses in LpEs secreted by *APOE2/E2* (LpE2), *APOE3/E3* (LpE3), and *APOE4/E4* (LpE4) pericytes. Points in graphs represent individual replicates (n = 3), bars represent the mean, and error bars ± SD. Pairwise comparisons were performed within each lipid subclass, *p < 0.05, **p < 0.01, ***p < 0.001, ****p < 0.0001 (adjusted p-values), ns — not significant.

**Supplementary Figure 4.**
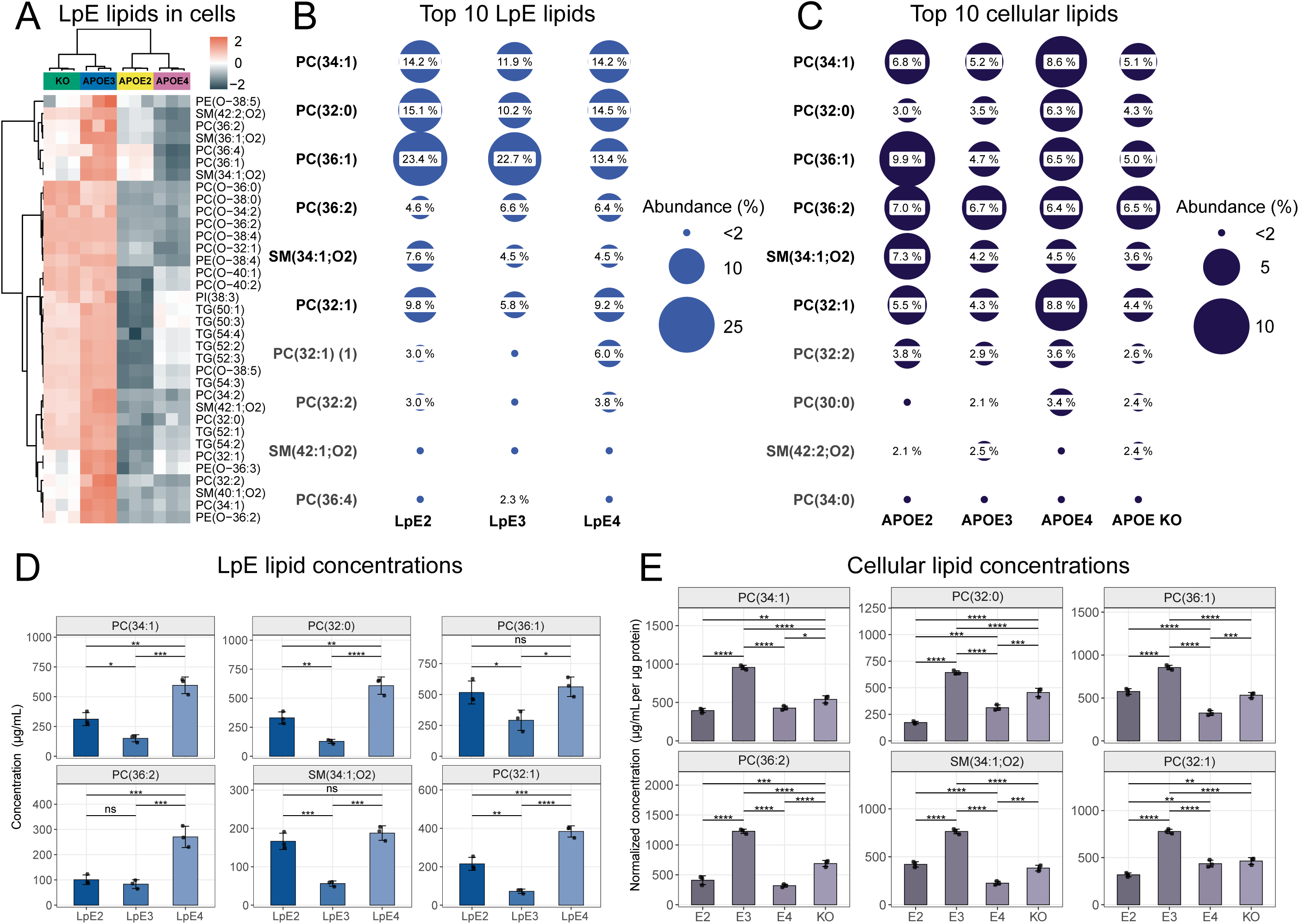
Cellular representation of lipids identified as LpE components in *APOE*-isogenic pericytes. A) Heatmap represents color-coded abundance (Z-scored, row-scaled) of cellular lipids in *APOE3/E3*, *APOE2/E2*, *APOE4/E4*, and *APOE* KO pericytes. Only lipids identified as pericyte-derived LpE components are shown. B) Dot plot shows top 10 lipids and their relative abundances in LpEs secreted by *APOE2/E2* (APOE2), *APOE3/E3* (APOE3), and *APOE4/E4* (APOE4) pericytes. Only relative abundances >2% are labeled. C) Dot plot shows top 10 cellular lipids and their relative abundances in *APOE2/E2* (APOE2), *APOE3/E3* (APOE3), *APOE4/E4* (APOE4), and *APOE* KO pericytes. Only relative abundances >2% are labeled. D) Absolute concentrations of most abundant lipid species in LpEs secreted by *APOE2/E2* (LpE2), *APOE3/E3* (LpE3), and *APOE4/E4* (LpE4) pericytes. E) Concentrations of most abundant cellular lipid species in *APOE2/E2* (E2), *APOE3/E3* (E3), *APOE4/E4* (E4), and *APOE* KO (KO) pericytes, normalized to total protein content. Points in graphs represent individual replicates (n = 3), bars represent the mean, and error bars ± SD. Pairwise comparisons were performed within each lipid subclass, *p < 0.05, **p < 0.01, ***p < 0.001, ****p < 0.0001 (adjusted p-values), ns — not significant. Only significant pairwise comparisons are shown when four groups are compared.

**Supplementary Figure 5.**
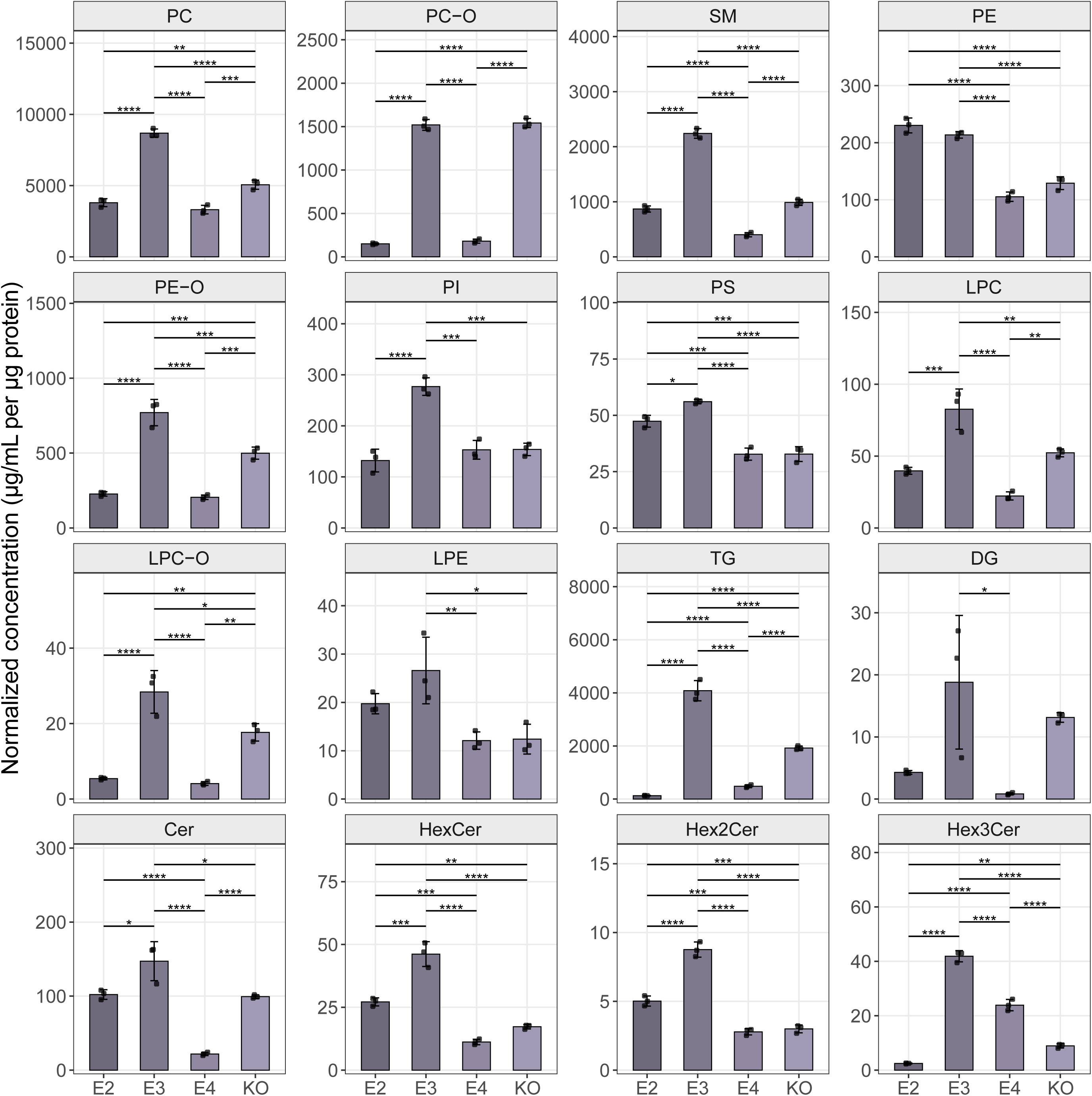
Lipid subclass concentrations in *APOE*-isogenic pericytes. Concentrations of cellular lipid subclasses in *APOE2/E2* (E2), *APOE3/E3* (E3), *APOE4/E4* (E4), and *APOE* KO (KO) pericytes, normalized to total protein content. Points in graphs represent individual replicates (n = 3), bars represent the mean, and error bars ± SD. Pairwise comparisons were performed within each lipid subclass, *p < 0.05, **p < 0.01, ***p < 0.001, ****p < 0.0001 (adjusted p-values). Only significant pairwise comparisons are shown.

**Supplementary Figure 6.**
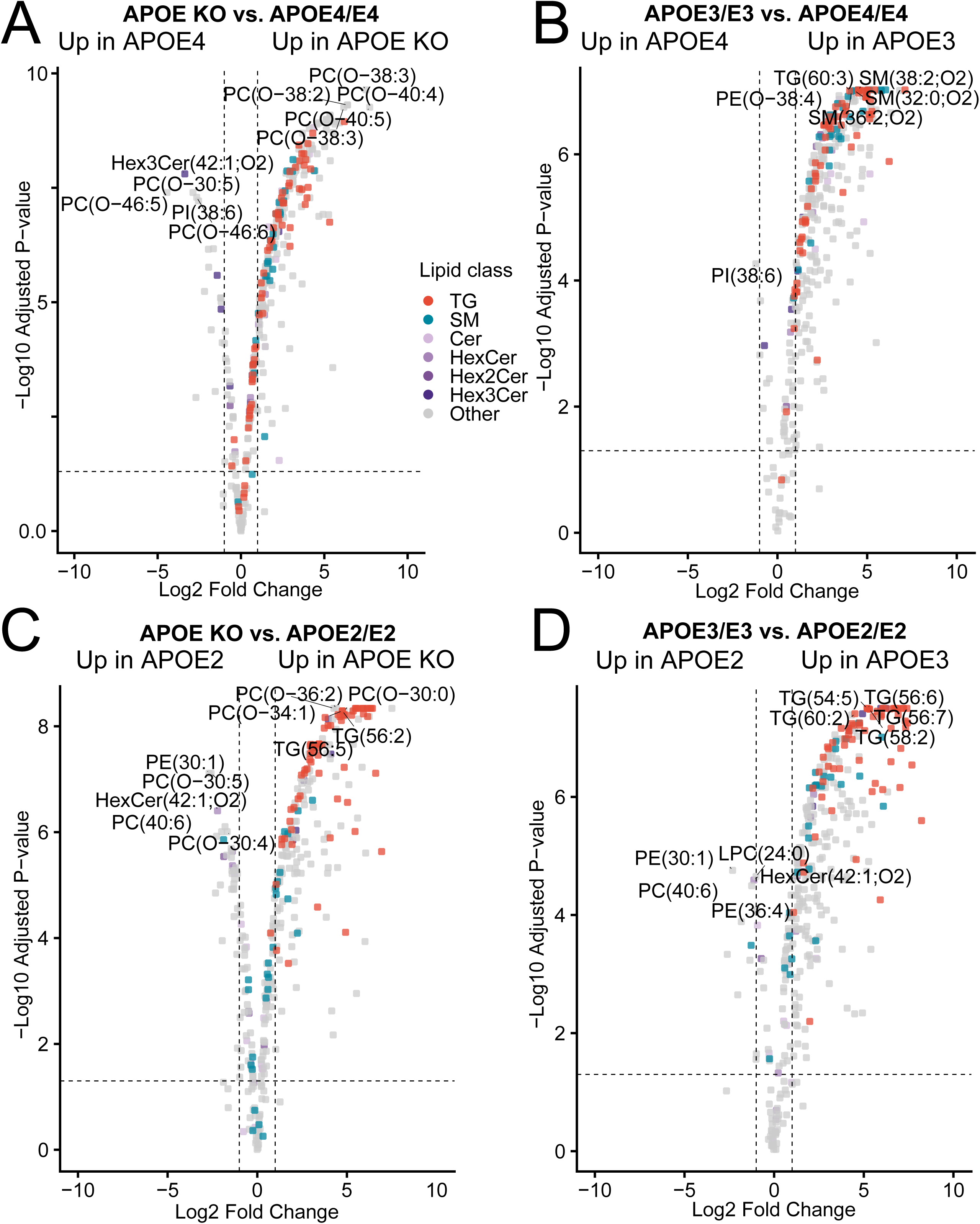
Differential expression analysis of cellular lipids. A-D) Differential expression of cellular lipids in *APOE* KO vs. *APOE4/E4* (A), *APOE3/E3* vs. *APOE4/E4* (B), *APOE* KO vs. *APOE2/E2* (C), and *APOE3/E3* vs. *APOE2/E2* (D) pericytes. Each point represents one lipid species. Significance thresholds: |log2FC| ≥ 1 (FC ≥ 2), FDR < 0.05. FC – fold change, FDR – false discovery rate.

**Supplementary Figure 7.**
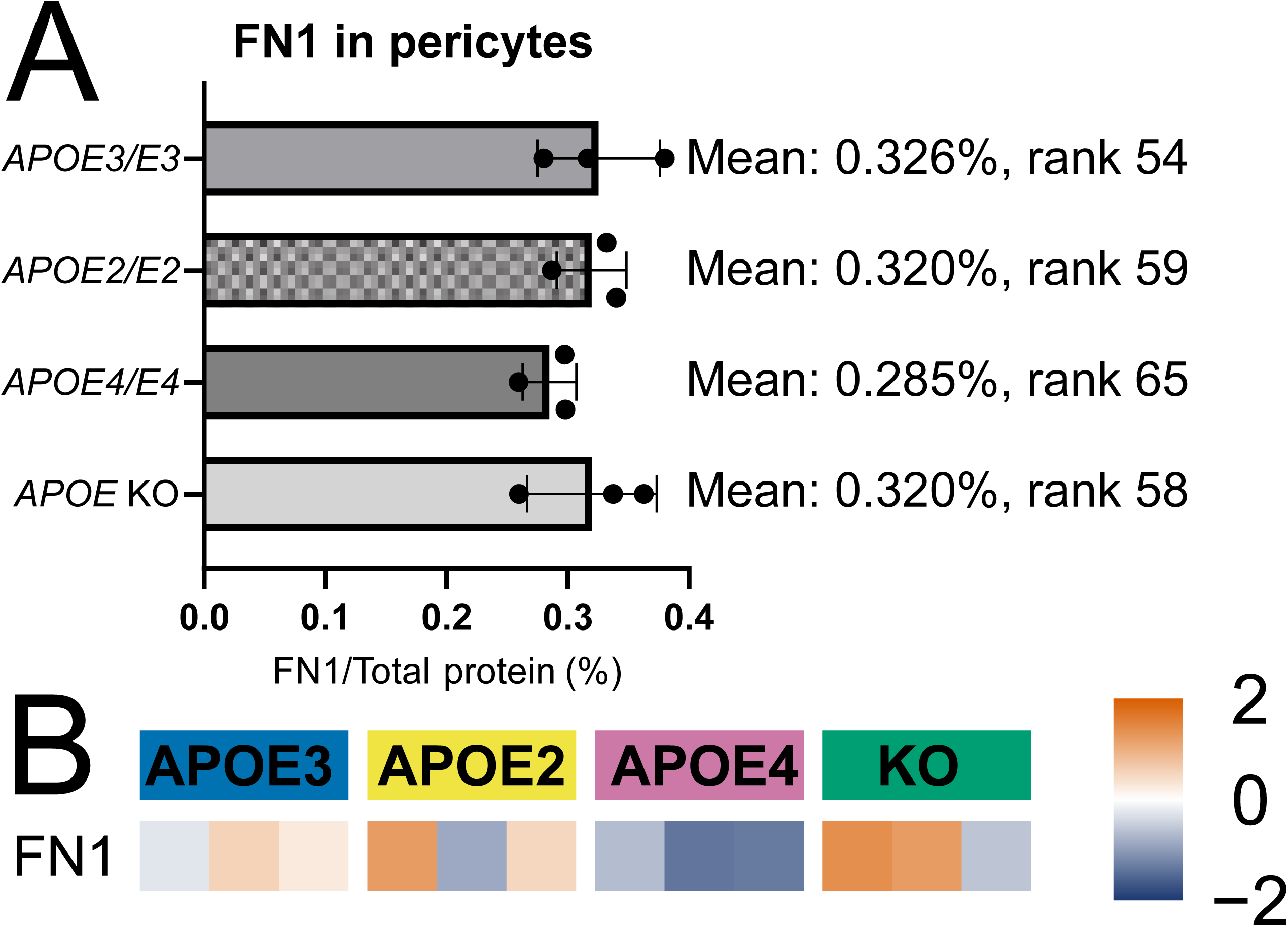
Cellular FN1 expression in *APOE*-isogenic pericytes. A) Cellular FN1 abundance in *APOE*-isogenic pericytes, calculated as percentage contribution of FN1 to total protein signal per sample. Points in graphs represent individual replicates (n = 3), bars represent the mean, and error bars ± SD. B) Heatmap represents color-coded abundance (Z-scored, row-scaled) of cellular FN1 in *APOE-*isogenic iPSC-derived pericytes.

## Supporting information

Sup. File 1. Protein filtering

Sup. File 2. Lipid filtering

Sup. File 3. Rel abundance LpE lipids

Sup. File 4. Rel abundance cellular lipids

Sup. File 5. Rel abundance LpE proteins

Sup. File 6. Rel abundance cellular proteins

Sup. Table 1. CSF LpE lipids comparison

Sup. Table 2. CSF LpE proteins comparison

